# Precise DNA Cloning via PAMless CRISPR-SpRYgests

**DOI:** 10.1101/2022.01.11.474553

**Authors:** Kathleen A. Christie, Jimmy A. Guo, Rachel A. Silverstein, Roman M. Doll, Megumu Mabuchi, Hannah E. Stutzman, Linyuan Ma, G. Brett Robb, Benjamin P. Kleinstiver

## Abstract

While restriction enzymes (REs) remain the gold-standard for manipulating DNA *in vitro,* they have notable drawbacks including a dependence on short binding motifs that constrain their ability to cleave DNA substrates. Here we overcome limitations of REs by developing an optimized molecular workflow that leverages the PAMless nature of a CRISPR-Cas enzyme named SpRY to cleave DNA at practically any sequence. Using SpRY for DNA digests (SpRYgests), we establish a method that permits the efficient cleavage of DNA substrates at any base pair. We demonstrate the effectiveness of SpRYgests using more than 130 gRNAs, illustrating the versatility of this approach to improve the precision of and simplify several cloning workflows, including those not possible with REs. We also optimize a rapid and simple one-pot gRNA synthesis protocol, which reduces cost and makes the overall SpRYgest workflow comparable to that of RE digests. Together, SpRYgests are straightforward to implement and can be utilized to improve a variety of DNA engineering applications.

---

Restriction enzymes (REs) transformed the field of molecular cloning by enabling and accelerating the assembly of recombinant DNA fragments^1^. REs commonly utilized for cloning applications recognize sequence motifs typically 4-8 base pairs in length and generate DNA double strand breaks (DSB) (**Fig. 1a**). Despite a diverse catalog of REs, there remain challenges for molecular cloning since the availability of these motifs in DNA substrates is stochastic, and to be useful, the RE motif should be conveniently located and generally must only occur once (so-called ‘single cutters’). Aside from purposefully designed multiple-cloning sites (MCSs) harboring several unique RE motifs (**Sup. Fig. 1a**), most RE motifs either do not occur in a substrate or are found more than once, preventing specific digestion of a DNA substrate to generate the desired fragments. Given these caveats, most laboratories purchase a repertoire of REs which can be costly and is incomplete since REs cannot comprehensively address all sequences.

**Fig. 1:**
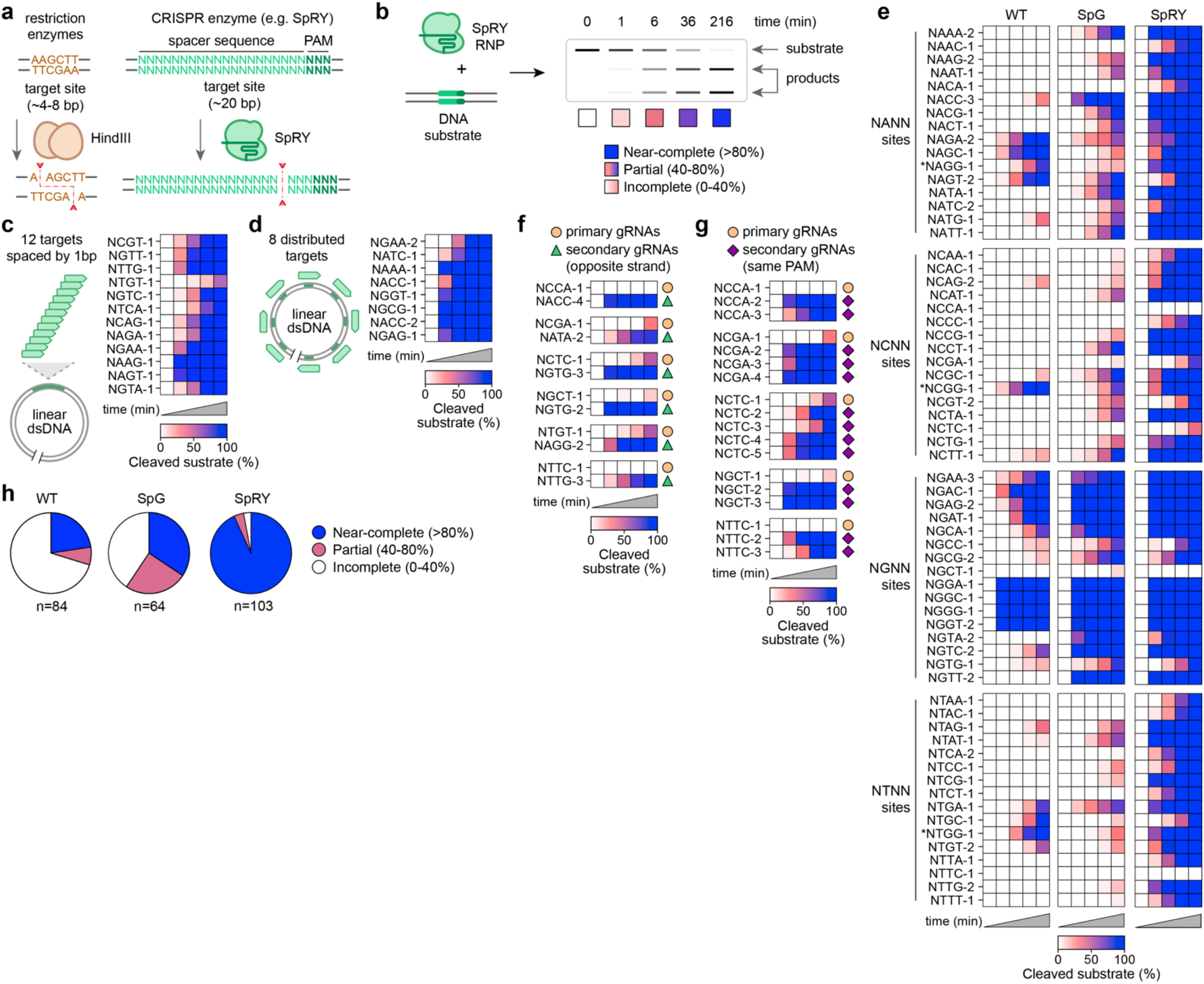
Characterization of SpRYgest in vitro cleavage efficiencies. (**a**) Comparison of restriction enzymes (REs) that require fixed 4-8 nt motifs and the near-PAMless Cas9 variant, named SpRY, that can target and cleave DNA substrates without sequence constraints. (**b**) Illustration of SpRYgest in vitro cleavage reaction workflow. Categorization of substrate cleavage is determined at the final timepoint, judged as near-complete (>80%), partial (40-80%), or incomplete (<40%) digests. (**c**, **d**) Initial SpRYgest experiments assessing the DNA cleavage efficiency of SpRY against a linearized plasmid substrate by targeting a specific region at 1 bp intervals and 8 gRNAs designed to be distributed across the substrate (panels **c** and **d**, respectively). (**e**) Comparison of the in vitro cleavage efficiencies of WT, SpG, and SpRY across 64 target sites representing all 2^nd^/3^rd^/4^th^ position combinations of an NNNN PAM. Sites with a shifted NNGG PAM are indicated with an asterisk. (**f**, **g**) SpRYgest results using additional secondary gRNAs for primary sites in **panels 1c** and **1e** for which partial or incomplete cleavage was observed. Secondary gRNAs were designed to target the opposite DNA strand placing the DSB at the same position as the primary gRNA (**panel f**), or to target different sites (spacers) but bearing the same PAMs as the primary gRNA (**panel g**). (**h**) Summary of the proportion of gRNAs that led to near-complete, partial, or incomplete substrate cleavage when using WT, SpG and SpRY. For **panels c-g**, cleavage of DNA substrates was quantified by capillary electrophoresis; mean shown for n = 3.

CRISPR-Cas nucleases are one potential alternative platform that can cleave DNA in a customizable manner. Most of the DNA-targeting specificity of Cas enzymes is provided by a guide RNA (gRNA) that can be programmed to bind to nearly any sequence^2^ (**Sup. Fig. 1b**). However, canonical Cas9 nucleases are also encumbered by the requirement to recognize a short DNA motif adjacent to the target site, known as the protospacer adjacent motif^3^ (PAM; **Sup. Fig. 1b**). The dependence on the availability of a PAM proximal to the target site prohibits precise cleavage of DNA substrates (**Sup. Fig. 1c**), including for *in vitro* DNA digests^4^. To overcome this limitation, a nearly PAMless CRISPR-Cas variant named SpRY was recently engineered^5^ that can target DNA sites with NNN PAMs in human cells, with a preference for NRN PAMs over NYN PAMs (where R is A or G and Y is C or T; **Fig. 1a** and **Sup. Fig. 1d**). Given that SpRY is no longer dependent on a PAM, we investigated whether SpRY could act as a fully programmable DNA endonuclease to cleave at any DNA base *in vitro,* vastly simplifying the design and execution of molecular cloning protocols.

We first investigated whether SpRY could generate DSBs along a DNA substrate at various locations harboring different PAMs. In our initial assays (**Fig. 1b**), we performed *in vitro* digests utilizing overexpressed SpRY protein from human cell lysates (**Sup. Figs. 2a** and **2b**) along with gRNAs produced using optimized and rapid *in vitro* transcription (IVT) conditions (**Sup Figs. 3a-3c** and **Sup. Note 1**). We assessed SpRY activity *in vitro* by performing SpRY DNA digests (SpRYgests) against 20 different target sites sampling NRN and NYN PAMs across a linearized plasmid substrate (**Figs. 1c** and **1d**). We utilized 12 gRNAs targeting a specific region at 1 bp intervals and 8 gRNAs distributed across the substrate (**Sup Fig. 4**). Between the two experiments, we observed near-complete cleavage of the substrate for 19 of 20 gRNAs, with the lone gRNA resulting in approximately 50% substrate digestion (**Figs. 1c**, **1d**). In comparison, wild-type (WT) SpCas9 digested only 4 of these 20 sites to near-completion (>80% digestion; **Sup Figs. 4c** and **4f**). These results provided evidence that SpRY could act as a potent PAM-agnostic endonuclease *in vitro.*

Intrigued by the PAMless nature of SpRYgests, we performed a large comparison of WT SpCas9, SpG (an SpCas9 variant previously engineered to target sites with NGN PAMs^5^), and SpRY using 64 additional gRNAs targeting a range of sites bearing all 2^nd^/3^rd^/4^th^ position combinations of an NNNN PAM (**Fig. 1e** and **Sup. Figs. 5a-c**). Similar to previous reports^6,7^, WT SpCas9 efficiently digested substrates when programmed with gRNAs targeting sites harboring NGG PAMs, and sometimes exhibited activity against sites with NAG, NGA, or shifted NNGG PAMs (**Fig. 1e** and **Sup. Fig. 5a**). SpG recapitulated its preference to edit substrates with NGN PAMs (**Fig. 1e** and **Sup. Fig. 5b**). Finally, although an NRN PAM preference was observed with SpRY in mammalian cells^5^, SpRY digested the substrate to near-completion when using 59 of 64 gRNAs, partially digested the substrate with 2 gRNAs, and exhibited low-to-no activity with the remaining 3 (**Fig. 1e** and **Sup. Fig. 5c**). Together, 93% (78/84) of the gRNAs initially used in these two sets of SpRYgests led to extensive cleavage.

Next, we investigated potential causes for incomplete substrate digestion with SpRY. First, for sites where the primary gRNA exhibited partial or incomplete cleavage, we tested the ability of a secondary gRNA targeted to the opposite strand to generate a DSB at the exact same location (**Fig. 1f**, **Sup. Figs. 6a** and **6b**). This strategy generates the same DSB but via a different target site and gRNA. We assessed the opposite-strand secondary gRNA approach for 6 primary gRNAs that did not reach >80% completion. Importantly, we observed >80% cleavage for 5 of 6 new secondary gRNAs targeted to the opposite strand and 78% cleavage for the 6^th^ gRNA (**Fig. 1f** and **Sup. Fig. 6b**), identifying a strategy to overcome low-activity gRNAs. Next, we explored whether low-activity sites could be attributed primarily to a spacer- or PAM-specific source. We tested additional gRNAs targeted to new sites/spacers bearing some of the PAMs that initially resulted in incomplete cleavage (**Fig. 1g** and **Sup. Fig. 6c**). For all 13 new gRNAs, we observed near-complete substrate cleavage, suggesting that the PAM is not a primary determinant for incompletely digested sites and that SpRY is generally PAMless *in vitro.* Collectively, our combined results using 103 gRNAs reveal the flexibility and effectiveness of SpRY for *in vitro* digests, with 93.2% of SpRYgests achieving near-complete substrate digestion, an efficiency dramatically higher than for WT SpCas9 or SpG^5^ (**Fig. 1h**).

The flexibility to generate DSBs at specific positions within plasmid substrates holds promise to simplify, accelerate, and improve the precision of molecular cloning applications. As an alternative to REs, another option aside from SpRYgest is the use of DNA-guided prokaryotic Argonaute (Ago) proteins^8,9^. To compare an Ago protein with SpRY, we initially examined the ability of commercially available *Thermus thermophilus* Argonaute (TtAgo) to generate custom DNA breaks when programmed with pairs of ssDNA guides (**Sup. Figs. 1e** and **7a**). In experiments adhering to the restrictive target site design considerations for TtAgo (**Sup. Fig. 7b** and **Sup. Note 2**) and despite performing metal ion and enzyme dose optimizations (**Sup. Figs. 7c** and **7d**, respectively),

TtAgo was only able to cleave 2 of 5 substrates to near-completion (**Fig. 2a** and **Sup. Fig. 7e**). By comparison, all ten SpRYgests targeting either strand for each of the five TtAgo sites reached near-complete cleavage (**Fig. 2a** and **Sup. Fig. 7e**). We then tested TtAgo against two positive control sites and the 20 sites that we initially examined with SpRY (where 19/20 resulted in near-complete digestion; **Figs. 1c** and **1d**). None of these 20 sites accommodated the restrictive TtAgo design requirements and we did not observe evidence of DNA cleavage at any of the 20 sites (**Fig. 2b** and **Sup. Fig. 7f**). Given that TtAgo fully digested the DNA substrate for only 8% of sites examined, at least under our current optimized conditions, TtAgo cannot generate DSBs *in vitro* as effectively as SpRY (**Fig. 2b**).

**Fig. 2:**
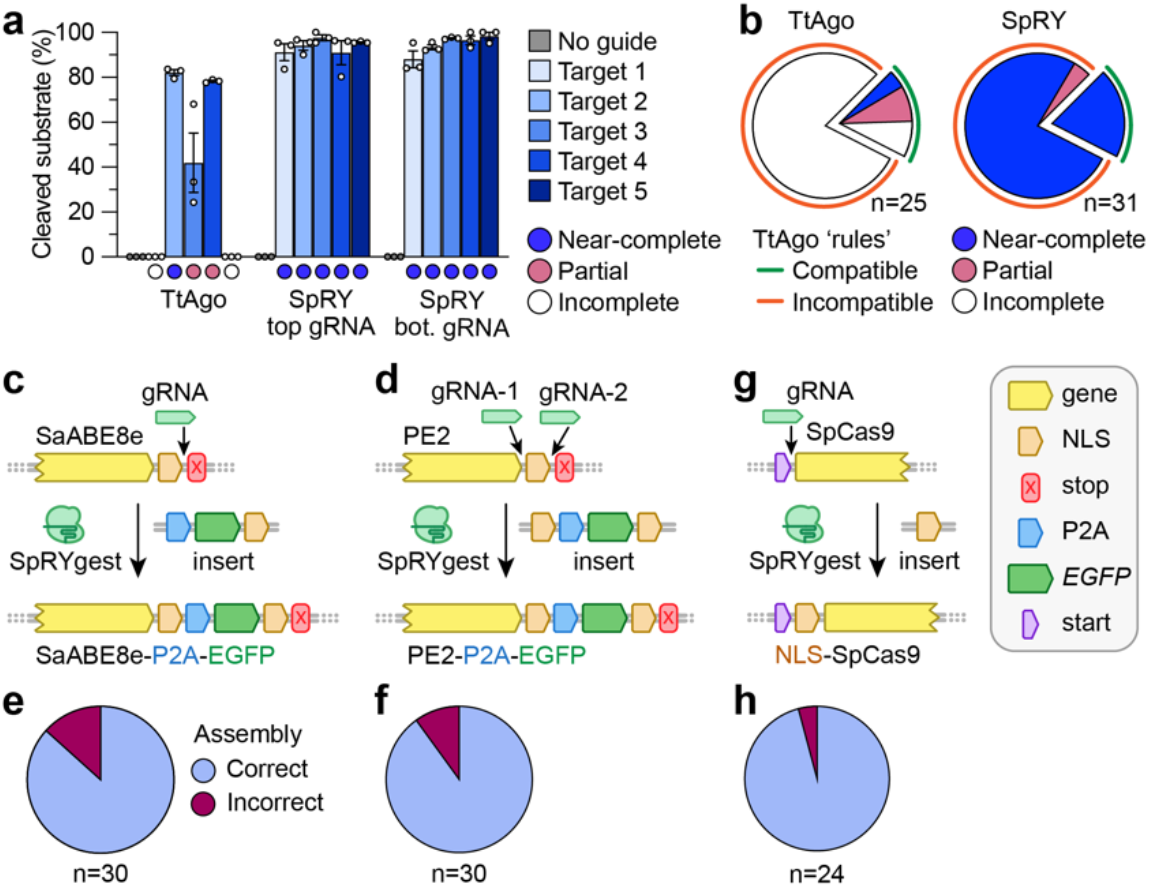
Molecular cloning via SpRYgest. (**a**) Comparison of the DNA cleavage efficiencies of TtAgo and SpRY across 5 sites designed to adhere to TtAgo guide requirements. TtAgo reactions were performed with pairs of 5’P-ssDNA guides. TtAgo and SpRYgest reactions were performed for 60 and 216 minutes, respectively. Individual datapoints, mean, and s.e.m. shown for n = 3. (**b**) Proportion of TtAgo and SpRY guides that led to nearly complete, partial, or incomplete cleavage on the 20 target sites from **Figs. 1c** and **1d** (see TtAgo results in **Supplementary Fig. 7f**) and the 5 sites from **Fig. 2a**. For **panels a** and **b**, cleavage of DNA substrates was quantified by capillary electrophoresis; mean shown for n = 3. (**c**, **d**) Schematics of the SpRYgest strategies to add P2A-EGFP sequences to SaCas9-ABE8e and PE2 via single and double SpRYgests, **panels c** and **d,** respectively. (**e**, **f**) Proportion of clones for which correct addition of the P2A-EGFP sequence was confirmed by Sanger sequencing for SaCas9-ABE8e and PE2 strategies, **panels e** and **f,** respectively. (**g**) Schematic of the SpRYgest strategy to add an NLS to the N-terminal end of SpCas9. (**h**) Proportion of clones for which correct addition of the N-terminal NLS was confirmed by Sanger sequencing.

To evaluate the practical utility of SpRYgests for molecular cloning applications, we first scaled up our cleavage reactions from ng to μg quantities of DNA substrate. Because these conditions necessitated the use large quantities of SpRY, we overexpressed and purified SpRY from *E. coli* (**Sup. Fig. 8**). We initially tested purified SpRY *in* vitro using 3 different gRNAs targeting sites with NAT, NCA, and NGG PAMs across 9 different temperatures. Our data revealed that, consistent with prior reports for WT SpCas9, reactions at 37 °C were optimal for SpRY (**Sup. Fig. 9a**). The cleavage efficiencies of these 3 gRNAs with purified SpRY were consistent with our previous results using SpRY from a human cell lysate (**Sup. Fig. 9b**). For sites that exhibited lower activities using the SpRY from lysate, SpRY protein often improved editing efficiency (**Sup. Fig. 9c**). Upon scaling up the cleavage reactions to μg amounts of DNA substrate, we sometimes observed non-specific degradation of the digestion products (**Sup. Fig. 10a**). Reactions with gRNA-only conditions led to non-specific nicking of the supercoiled plasmid substrate, suggesting carry-forward of DNase from template removal during the IVT reaction (**Sup. Figs. 10a** and **10b**). Omission of the of the DNase treatment during gRNA synthesis, or use of chemically synthesized gRNAs, eliminated the non-specific degradation of the SpRYgested DNA products (**Sup. Fig. 10c**).

We then explored the potential of SpRYgests to perform routine cloning applications where unique restriction sites were not available. First, we sought to precisely insert long ~1kb P2A-EGFP sequences into plasmids encoding two different genome editors, SaCas9-ABE8e^10^ and a prime editor^11^ (PE2) (**Figs. 2c** and **2d**, respectively). To do so, we performed single and double gRNA SpRYgests using 4 μg of supercoiled plasmid substrate (**Sup. Figs. 11a**-**11d**). For both reactions, complete SpRYgestion was observed. Conversely, no cleavage was observed when using TtAgo targeted to these sites (**Sup. Fig. 11e**). We then generated a PCR product encoding the P2A-EGFP sequence and cloned it into the digested plasmids via isothermal assembly^12^ (**Sup. Figs. 11a** and **11b**). Of the 30 resulting clones that we sequenced for each assembly, 26 and 27 were correct (**Figs. 2e** and **2f**). In another molecular cloning reaction, we performed a single SpRYgest of the N-terminus of a Cas9 expression plasmid to add a nuclear localization sequence (NLS) (**Fig. 2g** and **Sup. Fig. 12**). Isothermal assembly using a short PCR product was extremely efficient, leading to 23 out of 24 clones assembling correctly (**Fig. 2h**). Together, SpRYgests simplified these three practical and exemplary cloning experiments with assembly efficiencies comparable to typical RE-based digests.

Like genome editing experiments, off-target cleavage of closely related sequences could manifest in SpRYgests. When cloning the SaCas9-ABE8e-P2A-EGFP plasmid, we observed evidence of a very low-level secondary set of products likely caused by an off-target DSB (**Sup. Fig. 11c**). Closer inspection of the target site revealed that the off-target cleavage was the result of utilizing a gRNA that overlapped an NLS, for which there was a second NLS with high sequence homology elsewhere in the plasmid, bearing 3 mismatches (**Sup. Figs. 13a** and **13b**). We were able to completely mitigate off-target cleavage by utilizing a secondary gRNA targeted to the more unique sequence on the opposite strand (**Sup. Figs. 13c** and **13d**). Another potential method to eliminate off-target editing is to utilize a high-fidelity version of SpRY, SpRY-HF1^5,13^ (**Sup. Fig. 13e**), that has previously been shown to be more sensitive to mismatches. While SpRY-HF1 did not completely prevent off-target editing when utilizing the NLS-targeted gRNA at the final digestion timepoint (**Sup. Fig. 13f**), a comparison of SpRY and SpRY-HF1 when programmed with mismatched gRNAs revealed that in certain cases, SpRY-HF1 can reduce off-target cleavage *in vitro,* especially at earlier reaction timepoints (**Sup. Fig. 13g**).

Another potential application of performing a SpRYgest is to generate plasmid libraries bearing customized regions at any location rapidly and cost-effectively. As a proof-of-concept, we sought to investigate biological properties of SpCas9 by generating two saturation mutagenesis libraries with randomized nucleotides in regions of SpCas9 that are critical for either the catalytic activity^14^ (**Fig. 3a** and **Sup. Fig. 14a**) or the PAM preference^7,15^ of SpCas9 (**Fig. 3b** and **Sup. Fig. 14b**). We linearized an SpCas9-encoding plasmid via a double-SpRYgest and performed isothermal assembly reactions with ssDNA oligonucleotides encoding degenerate NNS codons (where ‘N’ is any nucleotide and ‘S’ is G or C). Via both Sanger and next-generation sequencing, we observed balanced representation of nucleotides in the modified positions (**Figs. 3c** and **3d**). The libraries also contained intentionally coded silent substitutions to enable assessment of library construction efficiency, which were introduced at > 99.8% suggesting highly effective synthesis with minimal background (**Figs. 3c** and **3d**).

**Fig. 3:**
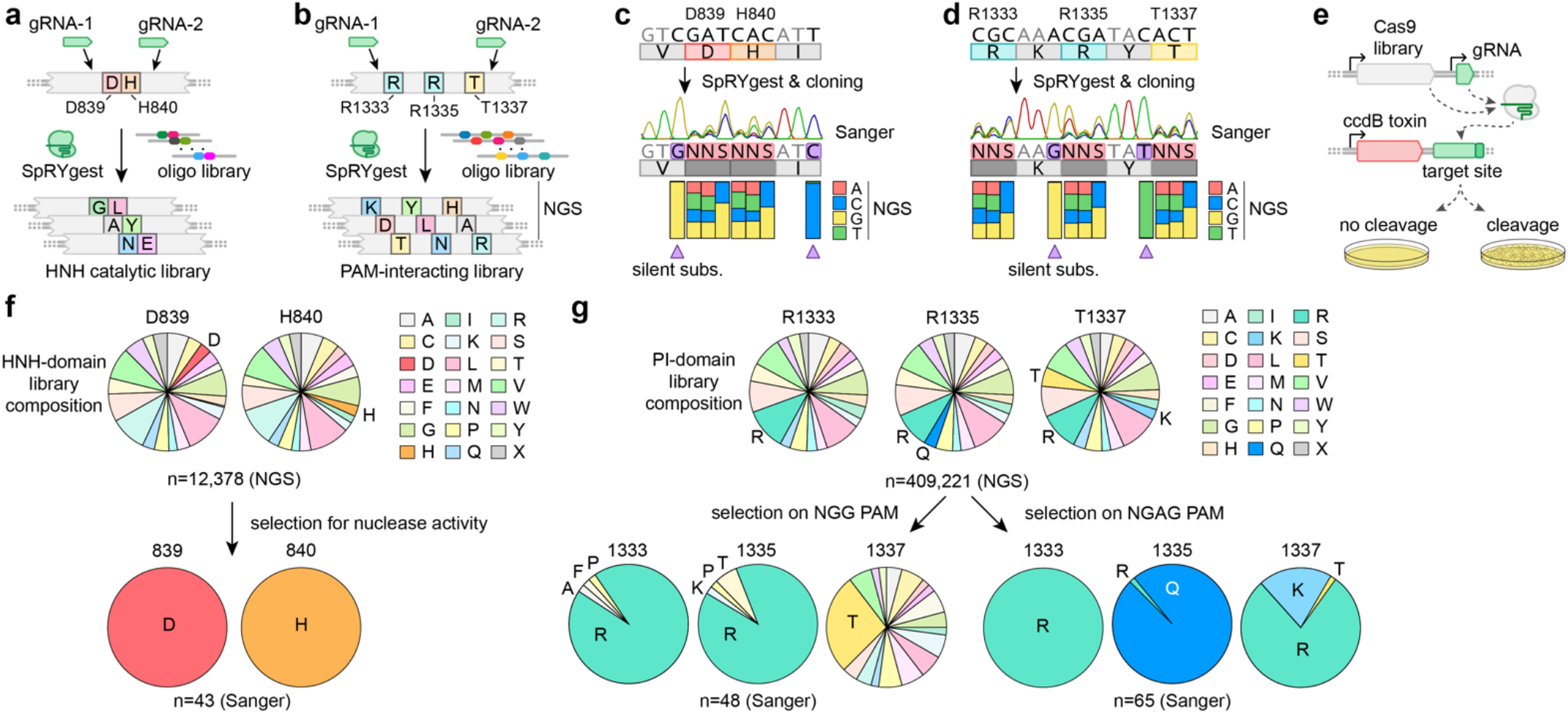
Generation of saturation mutagenesis libraries via SpRYgest. (**a,b**) Schematics of SpRYgest strategies to generate saturation mutagenesis libraries of SpCas9 residues important for catalytic activity (**panel a**) and NGG PAM preference (**panel b**). (**c,d**) Sanger sequencing traces and next-generation sequencing results from the libraries, illustrating the nucleotide diversity at mutated residues for the HNH-catalytic and PAM-interacting (PI) domain libraries, **panels c** and **d**, respectively. Recoded silent substitutions were intentionally included in the library to assess construction efficiency (highlighted in purple and indicated with a triangle). (**e**) Schematic of the bacterial positive selection assay^7,16,17^, which permits selection of cleavage competent SpCas9 enzymes from saturation mutagenesis libraries. Colonies survive on selective media only when SpCas9 and a gRNA cleave a target site on the toxic plasmid. Mutated regions of SpCas9 can be sequenced from the plasmids harbored within surviving colonies. (**f,g**) Post-selection results for cleavage competent SpCas9 variants from the catalytic domain HNH residue library (**panel f**), or from the PI domain library selected against toxic plasmids harboring target sites with NGG and NGAG PAMs (left and right sides of **panels g**, respectively). Pie charts illustrate the distribution of amino acids at each position in the pre-selection libraries (via NGS) and post-selection libraries (via Sanger sequencing of individual clones) in the top and bottom panels, respectively.

We then subjected these SpRYgest-constructed libraries to a previously described bacterial positive-selection^7,16,17^. Survival of colonies harboring transformed plasmids is dependent on the ability of SpCas9 to cleave a target site encoded within a selection plasmid that expresses a toxic gene (**Fig. 3e**). First, we performed experiments using the catalytic domain library that was varied at conserved HNH nuclease positions D839 and H840 (**Figs. 3a** and **3c**). Selection for cleavage-competent clones revealed that, as expected^18^, only SpCas9 variants encoding D839 and H840 were cleavage competent (**Fig. 3f**). We also evaluated the SpCas9 PAM-interacting (PI) domain library that was varied at amino acids critical for PAM recognition including R1333, R1335, and T1337 (**Fig. 3b** and **3d**). Selection of the library against a toxic plasmid encoding a target site with an NGG PAM led mostly to clones with R1333 and R1335, the two amino acid sidechains known to be important for specifying the guanines of the PAM^15^ (**Fig. 3g**). Interestingly, a variety of amino acids were observed at position 1337, though the native T1337 was enriched relative to the others. An additional selection with the same library against a target encoding an NGAG PAM led to enrichment of variants with R1333, R1335Q, and T1337R/K (**Fig. 3g**), consistent with expectations for amino acids that facilitate recognition of this non-canonical PAM^19,20^ (and previous results with the engineered variants SpCas9-VQR and SpCas9-VRQR^7,13^).

Finally, for SpRYgests to be more readily implemented, we sought to simplify and expedite the gRNA synthesis protocol (**Sup. Fig. 15a**). To do so, we experimented with various one-pot gRNA synthesis conditions and methods that require minimal hands-on time by combining the template generation and IVT steps (**Sup. Figs. 15a-15d** and **Sup. Note 3**). We identified conditions with shortened IVT reaction times (<4 hours) that generate high gRNA yields (**Sup. Figs. 15b** and **15c**). These optimized one-pot reaction conditions reduce SpRYgest hands-on times by approximately 3.5-fold, leading to workflows that are more similar to RE digests (**Fig. 4a**). Importantly, one-pot gRNA IVT reactions performed at two different scales for four different gRNAs all supported complete digestion of a plasmid substrate (**Fig. 4b**), reducing the enzymatic cost of gRNA synthesis by 300% to approximately $2.33 per gRNA (**Sup. Note 3**). The optimization of one-pot gRNA synthesis methods dramatically minimizes hands-on time, minimizes cost by scaling-down gRNA reactions, and makes the SpRYgest workflow more similar to other molecular cloning experiments (**Fig. 4a** and **Sup. Fig. 16**).

**Fig. 4:**
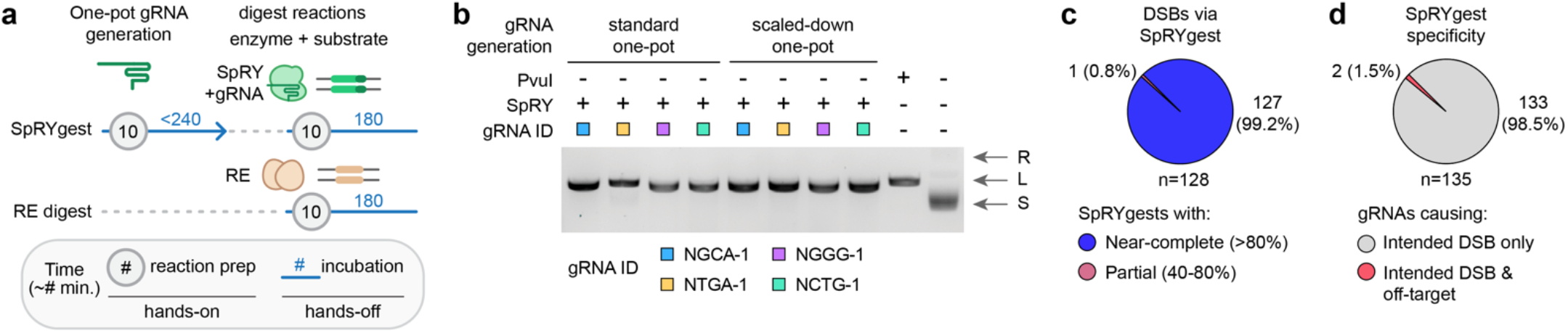
Optimization of rapid, efficient, and specific SpRYgest reactions. (**a**) Comparison of the hands-on and hands-off times of optimized SpRYgests (with gRNAs generated via one-pot reactions) versus RE digests. Approximate times in minutes are shown; lines not drawn to scale. The incubation time for one-pot gRNA generation can be shortened as needed (see **Sup. Fig. 15b**). (**b**) Agarose gel of SpRYgest reactions performed with gRNAs taken directly from standard or scaled-down one-pot gRNA generation reactions (20 μL or 5 μL one-pot reactions, respectively). Plasmid conformations are: R, relaxed; L, linear; S, supercoiled. (**c**) Proportion of unique DSBs that were successfully generated via SpRYgest in all experiments of this study. (**d**) Proportion of gRNAs for which off-target cleavage products were detected during SpRYgests with gRNAs used in all experiments of this study.

## Discussion

Here we establish a new paradigm of unconstrained DNA manipulation *in vitro* using a CRISPR-Cas enzyme, SpRY. We discover that SpRY is essentially PAMless when deployed in molecular cleavage reactions, and leverage this property for highly precise DNA digestion. With SpRYgests, standard and more complex cloning reactions are simplified, including the generation of saturation mutagenesis libraries to interrogate biological properties of impactful proteins (as demonstrated by our analysis of two SpCas9 domains). The PAMless nature of SpRY suggests that the NRN>NYN PAM preference we previously observed in human cells^5^ is less prominent under defined *in vitro* conditions. This is likely due to the fact that the ‘genome’ size of a plasmid is dramatically different than that of a eukaryotic organism, which influences SpRY target search time and how the preferred PAMs of a Cas variant are encountered and engaged^21–23^.

Across all experiments, we were able to generate a DNA break at every position that we sought to (**Fig. 4c**). For cases where the primary gRNA failed or was inefficient, reactions with the secondary gRNA that positions the DSB at the same position cleaved the substrate to near-completion (>80%) in all but one case ( where this single gRNA reached ~78% substrate digestion; **Fig. 4c**). We anticipate that the digestion efficiency can be further improved and expedited by simply increasing the amount of SpRY-gRNA complex included in the SpRYgest reaction, including for weaker gRNAs that don’t initially lead to complete cleavage. Our results also indicate that the substrate preferences of Ago proteins^8,9^ limit their applicability for DNA digests.

The unconstrained targeting range of SpRY eliminates the need for a large repertoire of REs. Even with a comprehensive catalog of REs, rarely does a single cutter RE site exist at the intended location of the DNA modification. SpRYgests require only a single source of the SpRY protein (versus dozens of REs) and a gRNA, which can be generated by IVT (**Fig. 4a** and **Sup. Fig. 15**). To streamline the IVT process, we optimized a rapid, simple, and affordable one-pot gRNA generation method, the product of which can be added directly to a SpRYgest reaction (**Fig. 4b** and **Sup. Note 3**). Despite the already minimal cost and rapid workflow of SpRYgest reactions, further optimization is possible (e.g. of the IVT reaction^24^). SpRYgest timelines are not substantially different than traditional cloning workflows, which are similarly dependent on the design and receipt of custom oligonucleotides (**Sup. Fig. 16**). We provide additional guidance on performing SpRYgests and considerations for experimental design (e.g. omitting the DNase treatment during IVT of gRNAs; see **Sup. Note 4**). Since Cas9 enzymes predominantly leave blunt DNA breaks, SpRYgests do not leave overhangs typical of most REs (a property largely obviated by the use of isothermal assembly^12^). However, paired SpRY nickases could in principle be utilized to generate cohesive ‘sticky’ ends as needed.

Over the course of this study when performing 136 separate SpRYgests, we observed only very low level off-target cleavage for 2 gRNAs (both of which were attributable to related sequences in the plasmids; **Fig. 4d** and **Sup Figs. 13b** and **13h**). These results indicate that achieving a single intended digestion product via SpRYgest is possible when considering other closely related sequences in the plasmid, and that simple methods can be used to mitigate off-target cleavage (by identifying when targeting the opposite strand would have fewer predicted off-target sites or by using SpRY-HF1, both of which can reduce or eliminate off-target cleavage). To prospectively identify gRNAs with closely matched off-target sites in DNA substrates, we have developed a web-based tool called SpOT-check (SpRYgest Off-Target checker; see **Sup. Note 5**).

Beyond a general usefulness for standard cloning, SpRYgests can improve various applications including saturation mutagenesis, domain minimization or shuffling^25^, depleting unwanted sequences from sequencing libraries^26^, target enrichment in sequencing protocols^27^, to generate more precisely terminated IVT templates, and other uses for the detection or diagnosis of infectious and genetic diseases. The flexibility of SpRYgests to generate DSBs at specific positions within DNA substrates holds promise to simplify, accelerate, and improve the precision of molecular applications, many of which not previously possible when using REs.

## Acknowledgements

We thank R.T. Walton for helpful suggestions and S. Mahendraker for assistance developing the web version of SpOT-check. K.A.C. is supported by a Massachusetts General Hospital (MGH) Fund for Medical Discovery (FMD) Fundamental Research Fellowship Award. B.P.K. acknowledges support from an MGH Executive Committee on Research Howard M. Goodman Fellowship.

## Author Contributions

K.A.C. and B.P.K. conceived of and designed the study. K.A.C., J.A.G., R.A.S., R.M.D., H.E.S. and L.M. performed experiments. M.M. and G.B.R. expressed and purified SpRY and SpRY-HF1. R.A.S. designed and wrote SpOT-check. All authors analyzed data. K.A.C. and B.P.K. wrote the manuscript draft and finalized the manuscript with input from all authors.

## Corresponding Author

Correspondence should be addressed to B.P. Kleinstiver

## Data Availability

All primary data will be made available as a Supplementary Table.

## Methods

### Plasmids and oligonucleotides

Descriptions and Addgene IDs for all plasmids used in this study are available in **Sup. Table 1**; new plasmids have been deposited with Addgene (https://www.addgene.org/Benjamin_Kleinstiver/). A list of all SpRYgest target sites is provided in **Sup. Table 2** that includes spacer sequences, PAMs, and gRNA generation methods. Oligonucleotide sequences and descriptions are available in **Sup. Table 3**. Target sites for TtAgo are listed in **Sup. Table 4**. Additional details for plasmids and oligonucleotides (oligos) are provided below in the respective sections. The SpOT-check computed off-target profiles for all gRNAs used in this study are available in **Sup. Table 5**.

### Human cell culture

Human HEK 293T cells (ATCC) were cultured in Dulbecco’s Modified Eagle Medium (DMEM) supplemented with 10% heat-inactivated FBS (HI-FBS) and 1% penicillin/streptomycin. The supernatant media from cell cultures was analyzed monthly for the presence of mycoplasma using MycoAlert PLUS (Lonza).

### Expression of and normalization of SpCas9-containing human cell lysates

Expression plasmids encoding WT SpCas9, SpG, and SpRY each with a -P2A-EGFP signal (RTW3027, RTW4177 and RTW4830, respectively) were used to generate human cell lysates containing SpCas9 proteins. Approximately 20-24 hours prior to transfection, 1.5×10^5^ HEK 293T cells were seeded in 24-well plates. Transfections containing 500 ng of human codon optimized nuclease expression plasmid and 1.5 μL TransIT-X2 were mixed in a total volume of 50 μL of Opti-MEM, incubated at room temperature for 15 minutes, and added to the cells. The lysate was harvested after 48 hours by discarding the media and resuspending the cells in 100 μL of gentle lysis buffer (containing 1X SIGMAFAST Protease Inhibitor Cocktail, EDTA-Free (Sigma), 20 mM Hepes pH 7.5, 100 mM KCl, 5 mM MgCl2, 5% glycerol, 1 mM DTT, and 0.1% Triton X-100). The amount of SpCas9 protein was approximated from the whole-cell lysate based on EGFP fluorescence. SpCas9 lysates were normalized to 180 nM fluorescien (Sigma) based on a standard curve. Fluorescence was measured in 384-well plates on a DTX 880 Multimode Plate Reader (Beckman Coulter) with λ_ex_ = 485 nm and λ_em_= 535 nm.

### Production of gRNAs

The DNA substrates required to transcribe gRNAs were generated via two methods. First, plasmids for IVT of SpCas9 gRNAs were generated by annealing and ligating duplexed oligos (see **Sup. Table 3**) corresponding to spacer sequences into BsaI-digested MSP3485 for T7 promoter-driven transcription of gRNAs. The derivative pT7-spacer-gRNA plasmids were digested with HindIII (NEB) to permit run-off transcription near the 3’ end of the SpCas9 gRNA. Secondly, oligo-derived DNA templates for IVT were generated by combining a target specific oligo (encoding a T7 promoter, spacer sequence, and partial sequence of the SpCas9 crRNA) and a common SpCas9 gRNA scaffold oligo (oKAC682), and then incubating with either Klenow Fragment (3’→5’ exo-) (New England Biolabs (NEB), M0212L) in 1x NEBuffer 2 at 37 °C for 30 minutes, or Q5 polymerase (NEB) using the following program: 2 minutes 98 °C; 5 cycles of (10 seconds 98 °C, 10 seconds 65 °C, 30 seconds 72 °C); 5 minutes 72 °C. Plasmid or oligo-derived transcription templates were cleaned up using a MinElute PCR Purification Kit (Qiagen). SpCas9 gRNAs were transcribed at 37 °C for 16 hours using the T7 RiboMAX Express Large Scale RNA Production Kit (Promega). For gRNAs utilized in *in vitro* cleavage reactions containing SpRY from human cell lysates, the 37 °C incubation was followed by the addition of 1 μL RQ1 DNase at 37 °C for 15 minutes to degrade the DNA template. The DNase treatment step was omitted when preparing most gRNAs utilized for scaled-up SpRYgest reactions with purified SpRY protein. Following transcription and optional DNase treatment, gRNAs were purified using paramagnetic beads (prepared as previously described^28^; GE Healthcare Sera-Mag SpeedBeads from Fisher Scientific, washed in 0.1X TE and suspended in 20% PEG-8000 (w/v), 1.5 M NaCl, 10 mM Tris-HCl pH 8, 1 mM EDTA pH 8 and 0.05% Tween20) and refolded by heating to 90 °C for 5 minutes and then cooling to room temperature at 1 °C every 2 seconds. Synthetic gRNAs were purchased from Synthego.

For one-pot gRNA IVT reactions, we utilized two general methods. First, gRNAs were generated using the EnGen sgRNA Synthesis Kit (NEB, E3322S) according to the manufacturer recommended protocol, or the EnGen sgRNA Synthesis Kit with increased oligo concentrations (final concentrations of 0.75 μM target-specific oligo and 0.75 μM common SpCas9 gRNA scaffold oligo (oKAC682)). The DNase step was omitted. Second, we also optimized a separate one-pot gRNA synthesis method using other commercial reagents. In this second method, 20 μL one-pot reactions were assembled containing final amounts or concentrations of 2.5 U Klenow Fragment (3’→5’ exo-), target-specific oligo at 0.5 or 1.5 μM (for standard or scaled-up reactions, respectively), common SpCas9 gRNA scaffold oligonucleotide (oKAC682) at 0.25 or 0.75 μM (for standard or scaled-up reactions, respectively), 125 μM dNTPs, 1x RiboMAX Express T7 Buffer (Promega, P1320), and 2 μL T7 Express Enzyme Mix (Promega, P1320) and incubated at 37 °C for 4 hours unless otherwise indicated. Appropriately scaled 5 μL reactions were assembled for smaller-scale one-pot reactions. For one-pot gRNAs used in SpRYgest reactions, the Promega recommended RQ1 DNase treatment was omitted and no clean-up of the gRNA was performed. To quantify gRNA yield, separate IVT reactions were performed that included the RQ1 DNase step and were purified using paramagnetic beads. Note that gRNA yield will vary based on incubation time.

### Expression and purification of SpRY and SpRY-HF1 proteins

*E. coli* codon optimized SpRY and SpRY-HF1 coding sequences including an N-terminal MKIEE tag and C-terminal SV40 NLS and 6x histidine tag were synthesized (GenScript, NJ, USA) and cloned into pET28 expression vectors. The SpRY and SpRY-HF1 expression constructs and were used to express and purify the proteins as described previously^29^. Briefly, *E. coli* strain NiCo21(DE3) (C2529H from NEB) harbouring the recombinant construct was grown in 1-2 L of LB medium with 40 μg/mL Kanamycin at 30°C until mid-log phase. Overexpression of the target protein was induced by adding IPTG to a final concentration of 0.4mM with shaking overnight at 18°C. Cells were harvested and target protein expression was assessed by SDS PAGE prior to purification. Cells were disrupted by sonication in breakage buffer (50mM Tris-HCl (pH8.0), 300mM NaCl, 1mM EDTA, 1mM DTT, 2% (v/v) glycerol) supplemented with PMSF. The supernatant was passed through HiTrap DEAE Sepharose (Cytiva, MA, USA) in column buffer (20mM Tris(pH7.5) and 250mM NaCl) followed by subsequent purification on a HisTrap HP column (Cytiva). After 16x column volume wash in buffer containing 20mM T ris pH7.5, 250mM NaCl, 40mM imidazole, target proteins were eluted using a 40mM to 750mM imidazole gradient in the same buffer. Pooled fractions containing the proteins were further purified by loading onto HiTrap heparin HP columns (Cytiva), washed with 6 column volumes of a buffer containing 20mM Tris (pH8.0), 1mM EDTA, and 1mM DTT, and eluted using a 0.25 to 2M NaCl gradient in the same buffer. Pooled fractions were dialyzed in SEC column buffer (20mM HEPES (pH8.0), 250mM KCl, and 1mM DTT) and concentrated using an Amicon^®^ Ultra-15 Centrifugal Filter Unit with 100 kDa molecular weight cut-off. Concentrated fractions were loaded on to a HiLoad 16/600 Superdex 200 pg column (Cytiva) using a 1 mL sample loop. Size exclusion chromatography was performed in SEC column buffer with a flow rate of 0.5mL/min. Eluted fractions were assessed by SDS-PAGE, pooled, dialyzed in storage buffer (20mM Tris (pH7.5), 300mM NaCl, 0.1mM EDTA, 1mM DTT and 50% (v/v) glycerol), and stored at −20°C. Protein concentration was determined by Bradford assay using BSA for standards.

### *In vitro* cleavage reactions using SpCas9 from lysates

Plasmid KAC833 linearized with HindIII (NEB) was used as the DNA substrate for most *in vitro* cleavage reactions unless otherwise stated. SpCas9 ribonucleoprotein (RNP) complexes were formed by mixing 9 μL of SpCas9-containing normalized whole-cell lysate (normalized to 180 nM Fluorescein) with 11.25 pmol of transcribed or synthetic gRNA, and incubating for 5 minutes at 37 °C. Cleavage reactions were initiated by the addition of 34.82 fmol of linearized plasmid (digested with HindIII (NEB)) and buffer to bring the total reaction volume to 22.5 μL with a final composition of 10 mM HEPES pH 7.5, 150 mM NaCl, and 5 mM MgCl2. Reactions were performed at 37 °C and aliquots were terminated at timepoints of 1,6, 36 and 216 minutes by removing 5 μL aliquots, mixing with 5 μL of stop buffer (50 mM EDTA and 2 mg/ml Proteinase K (NEB)), and incubating at room temperature for 10-minutes. Cleavage fragments were purified using paramagnetic beads and quantified via QIAxcel capillary electrophoresis (Qiagen). The relative abundances of substrate and products were analyzed using QIAxcel ScreenGel Software (v1.5.0.16, Qiagen) and plotted using GraphPad Prism 9 (v9.2.0).

### *In vitro* cleavage reactions using purified SpRY

Small-scale *in vitro* cleavage reactions were performed as described above, except using 0.6-1 μM purified SpRY per reaction pool instead of 9 μL of SpCas9-containing normalized whole-cell lysate (0.6 μM in **Sup. Figs. 9a** and **9b** and 1 μM in **9c**). For scaled-up digests, 4 μg of supercoiled plasmid DNA was incubated at 37 °C for 3 hours with purified SpRY protein at a final concentration of 1 μM and IVT gRNA (prepared without DNase treatment) at a final concentration of 2 μM in Buffer 3.1 (NEB). Reactions were stopped by the addition of 1 μL of Proteinase K (NEB) and incubated at room temperature for 15 minutes. Cleavage fragments were resolved by 0.8% agarose gel electrophoresis with 1 μL of 1 kb Plus DNA Ladder (NEB) and visualized by ethidium bromide staining. Digestion products were purified using a QIAquick Gel Extraction Kit (Qiagen).

### Molecular cloning reactions using purified SpRY

The C-terminal P2A-EGFP sequence was added to SaABE8e or pCMV-PE2 (Addgene IDs 138500 and 132775, respectively), and the N-terminal BPNLS was added to an SpCas9 plasmid similar to pCMV-T7-SpCas9 (Addgene plasmid ID 139987) via isothermal assembly. Reactions contained approximately 5 μL of isothermal assembly mix (prepared similar to as previously described^12^) or NEBuilder HiFi (NEB), 0.01 pmol of plasmid linearized via SpRYgest, and 0.03 pmol of PCR product insert in a final volume of 10 μL, and incubated at 50°C for 60 minutes. Cloning reactions were transformed into chemically competent XL1-Blue *E. coli* cells and grown at 37 °C for approximately 16 hours. Individual colonies were grown overnight at 37 °C, miniprepped (Qiagen), and fidelity of cloning was verified via Sanger sequencing. Saturation mutagenesis plasmid libraries for were constructed by incubating 0.02 pmol of BPK848 (linearized via SpRYgest) with 0.1 pmol of ~60bp ssDNA oligo (either oBK9102 or oBK9104) with 10 μL NEBuilder HiFi DNA Assembly Master Mix (NEB) in a final volume of 20 μL, and incubated at 50 °C for 15 minutes. NEBuilder reactions were cleaned up via MinElute (Qiagen) and eluted in 10 μL water, transformed into 100 μL of electrocompetent XL1-Blue *E. coli,* and recovered in 3 mL of SOC for 1 h at 37 °C. Next, 2 μL of the transformation recovery media was plated on LB + chloramphenicol, where the number of colonies following overnight at 37 °C were used to estimate library complexity. The remaining recovery was grown overnight in 150 mL LB + chloramphenicol and plasmid DNA was isolated by MaxiPrep (Qiagen). The complexity of the SpCas9 catalytic and PAM domain libraries were estimated to be 77,400 and 292,600 respectively. Plasmid libraries were sequenced via Sanger sequencing and NGS. For NGS, PCR amplicons were generated from the plasmids using primer pairs oKAC1589/oKAC1590 (for the catalytic domain) or oKAC1591/oKAC1592 (for the PI domain) and sequenced on a MiSeq (Illumina) to a depth of 12,378 and 409,221 reads for the catalytic and PI domain libraries, respectively. The resulting data was analyzed using CRISPResso2^30^ to generate allele tables.

### *In vitro* cleavage reactions using TtAgo

For TtAgo reactions, 5’-phosphorylated DNA guides were either purchased from Integrated DNA technologies or generated by incubating an unmodified oligonucleotide with T4 Polynucleotide Kinase (NEB) at 37 °C for 30 minutes, followed by heat-activation at 65°C for 20 min. Complexes of TtAgo programmed with ssDNA guides were prepared by combining final concentrations of 1 pmol TtAgo (NEB) and 2 pmol 5’-phosphorylated ssDNA guides and incubating at 70 °C for 20 minutes. Cleavage reactions were performed by combining TtAgo complexes with either 79.85 fmol of linearized KAC833 plasmid substrate (digested with PvuI, NEB) or 79.85 fmol supercoiled plasmid DNA (KAC1151 or MNW95) in ThermoPol buffer (NEB) with a final concentration of 10mM MgSO4. Reactions were performed at 80 °C for 60 minutes and terminated by the addition of 1 μL of Proteinase K (NEB). Cleavage fragments from pre-linearized substrates were purified using paramagnetic beads and quantified and analyzed as described above. Cleavage fragments from scaled-up plasmid DNA digests were resolved by 0.8% agarose gel electrophoresis and visualized by ethidium bromide staining.

### Bacterial-based positive selection assay

Target plasmids for the selection assays were generated by cloning duplexed oligonucleotides into XbaI and SphI-digested p11-lacY-wtx1 (Addgene ID 69056)^16^ as previously described^7^, which contains an arabinose-inducible ccdB toxin gene. The derivative toxin-expressing plasmids contain target sites harboring either NGG or NGA PAMs (BPK740 and BPK754, respectively). To perform the selections, electrocompetent *E. coli* BW25141(λDE3)^17^ containing a toxin-expressing plasmid were transformed with BPK848-derived plasmids that express the SpCas9 variant libraries (encoding randomized codons in specified positions) in addition to a gRNA, both from separate T7 promoters. Following a 60-minute recovery in SOC media, transformations were spread on LB plates containing either chloramphenicol and 10 mM dextrose (non-selective) or chloramphenicol + 10 mM arabinose (selective). Transformation efficiency was assessed based on colony count from non-selective plates. The catalytic library selection resulted in approximately 9e4 colonies (from sampling approximately 87x library coverage). The PI domain library selections for NGG PAMs and NGA PAMs resulted in approximately 6e5 and 6.4e4 colonies (from sampling approximately 18x and 2x coverage of the libraries, respectively). Surviving colonies from selective plates were picked as single colonies for miniprep (Qiagen) followed by Sanger sequencing to verify the identities of the mutated amino acids.

