## Supplementary Materials for "Precise DNA Cloning via PAMless CRISPR-SpRYgests"

#### Supplementary Material

##### Supplementary Tables

**Supplementary Table 1**: List of plasmids used in this study

**Supplementary Table 2**: List of SpRYgest target sites in this study

**Supplementary Table 3**: List of oligonucleotides used in this study

**Supplementary Table 4**: List of TtAgo target sites used in this study and their characteristics

**Supplementary Table 5**: SpOT-check results for SpRYgest sites in this study

***Note:*** *All Supplementary Tables are attached together as a single .xlsx file, separated by tabs.*

##### Supplementary Notes

**Supplementary Note 1:** Rapid synthesis of gRNAs by oligo extension and IVT p.2

**Supplementary Note 2:** Target site and guide design considerations for TtAgo p.2

**Supplementary Note 3:** Optimization of one-pot gRNA IVT reactions p.3

**Supplementary Note 4:** Design considerations and recommendations for SpRYgests p.4

**Supplementary Note 5:** Annotation of substrate off-targets via SpOT-check p.5

##### Supplementary Figures

Supplementary Figures 1-16 p.6-27

##### Supplementary References

Supplementary References p.28

### Supplementary Notes:

#### Supplementary Note 1: Rapid synthesis of gRNAs by oligo extension and IVT

To expedite the production of gRNAs for custom SpRYgests, we compared several methods to generate a dsDNA template for IVT of gRNAs. First, we cloned duplexed oligonucleotides (oligos) corresponding to the spacer sequence into a BsaI-digested entry plasmid that harbors a T7 promoter and the SpCas9 gRNA scaffold (MSP3485). The resulting pT7-spacer-gRNA plasmid can be linearized using HindIII for run-off transcription that terminates several bases after the 3' end of the gRNA. Secondly, we generated DNA substrates encoding pT7-gRNAs via oligo extension using either Klenow Fragment (3'→5' exo-) or Q5 polymerase. Oligo extension is performed by incubating a target-specific oligo encoding the T7 promoter, the 20bp spacer and a portion of the SpCas9 scaffold, where the latter section permits annealing with a common bottom-strand oligo (oKAC682) encoding the remainder of the SpCas9 scaffold. The common oligo partially anneals to the target-specific oligo, allowing oligo extension to generate a full-length dsDNA template for IVT (**Sup. Fig. 3a**). We observed that the three gRNA synthesis methods led to gRNAs that exhibited comparable cleavage efficiencies during SpRYgests (**Sup. Fig. 3b**). We further optimized this process by combining the dsDNA template generation and IVT steps (**Fig. 4a** and **Sup. Fig. 15a**). We demonstrated that the gRNA yield from one-pot reactions performed for as little as one hour generate sufficient gRNA quantities for full-scale SpRYgests (**Sup. Fig. 15**). The one-pot gRNA synthesis method only involves approximately 10 minutes of hands-on time and incubations between 1 to 4 hours (**Sup. Figs. 15a**-**15c 16a**). This timing is in comparison to generating a gRNA using a cloned IVT template plasmid by RE digest and ligation, which can take at least 3 days. For each of these methods, gRNA synthesis can be performed in parallel to ordering PCR primers that are otherwise required for cloning, making the timing of SpRYgests not substantially different than when using REs (**Sup. Fig. 16**).

#### Supplementary Note 2: Target site and guide design considerations for TtAgo

It was previously demonstrated that *Thermus thermophilus* Argonaute (TtAgo) can be directed to generate a DSB in a dsDNA template using a pair of 5'-phosphorylated ssDNA guides and that it has a strong preference for AT-rich regions of substrates[1]. New England Biolabs has also characterized TtAgo guide requirements: https://www.neb.com/tools-and-resources/usage-guidelines/guidelines-for-optimization-of-tth-argonaute-ttago-reactions. Their recommendations for TtAgo guide design include: 1) to design ssDNA guides with a 5' matched thymidine, 2) to avoid adenosine in the 12$^{th}$ position of guides, 3) to select target sites with low GC content, 4) that the addition of ET SSB may aid in cutting higher GC content substrates under specific conditions, and 5) ThermoPol Reaction Buffer with 2mM magnesium(II) is optimal for most reactions.

All TtAgo reactions were conducted following the protocol outlined by NEB (Protocol for generating break in phosphodiester backbone with Tth Argonaute (TtAgo, NEB #M0665)). As a positive control we ordered a 5'-phosphorylated guide pair from Swarts *et al.*[1] (oKAC1347 and oKAC1348) and cloned 98 base pairs of the 17% GC content target region into pUC19. We then designed 5 pairs of ssDNA guides following the guidelines outlined above. While we found the positive control TtAgo guide could efficiently introduce a DSB in the linearized target plasmid (**Sup. Fig. 10e**), otherwise, TtAgo was only able to cleave 2 of 5 sites to near-completion when following NEB guide design rules (**Fig. 2a** and **Sup. Fig. 10d**). We found that increasing the $MgSO_4$ concentration from 2mM to 10mM slightly improved cleavage efficiency (**Sup. Fig. 10b**), however the increase in magnesium(II) concentration did not rescue activity on failed sites.

**Supplementary Note 3: Optimization of a one-pot gRNA IVT reaction**

To streamline the process of performing a SpRYgest, we sought to optimize a one-pot gRNA synthesis reaction that could simultaneously produce the dsDNA *in vitro* transcription template and *in vitro* transcribe the gRNA (**Sup. Fig. 15a**). The one-pot gRNA product could then be directly combined with SpRY protein and the DNA substrate to assemble the complete SpRYgest reaction (**Fig. 4a**).

To explore the feasibility of a one-pot gRNA synthesis method, first we determined the gRNA yields at various reaction times for our previously established gRNA synthesis method (**Supplementary Note 1**) and a commercially available one-pot kit (EnGen sgRNA Synthesis Kit; NEB, # E3322S) (**Sup. Fig. 15b**). While our standard IVT method and the one-pot EnGen kit generated sufficient quantities of RNA by 4 hours for two different gRNAs, the standard IVT method yielded approximately 4-fold more gRNA at each timepoint (**Sup. Fig. 15b**). We then determined whether we could assemble a more efficient one-pot reaction that could generate comparable gRNA yields to our standard IVT method. To do so, we examined seven different combinations of reagents, including one-pot reactions comprised of Klenow Fragment (3'→5' exo-) (NEB, M0212L) to generate the IVT template, and RiboMAX T7 Express Enzyme Mix (Promega) to transcribe the gRNA (**Sup. Fig. 15c**). Other variations included the additional of 3x more target-specific and common oligos (for both our homebrew one-pot and the EnGen kit), or scaling down the reagents used by our homebrew one-pot method by 1/4 to reduce cost. Of all combinations tested, we found that our new one-pot and scaled-down one-pot reactions consistently generated greater quantities compared to our previous standard 2-step IVT method or the EnGen kit (**Sup. Fig. 15c**).

With each of the seven gRNA synthesis methods that we assessed for four different gRNAs, we then performed SpRYgest reactions using the unpurified and unquantified gRNA transcription products directly in SpRYgest reactions containing 859.5 fmol of plasmid substrate (**Sup. Fig. 15d**). For 3 of 4 gRNAs, we observed complete cleavage for all seven gRNA synthesis methods. However, for the remaining gRNA (site NTGA-1), the homebrew one-pot and EnGen reactions containing 1x target-specific and common oligos failed to reach completion (with homebrew one-pot outperforming EnGen; **Sup. Fig. 15d**). The addition of 3x both the target-specific and common oligos to either method rescued gRNA yield to sufficient quantities to permit SpRYgest reactions proceeding to completion. Furthermore, to identify methods that make gRNA IVT more cost-effective, we also tested scaling down the reagents used by our homebrew one-pot method by 1/4. The scaled-down one-pot reactions generated comparable yields for the four gRNAs per μL of transcription reaction relative to the full-scale one-pot (**Sup. Fig. 15c**), and was also effective when used directly in SpRYgest reactions resulting in complete substrate digestion (**Fig. 4b**).

Thus, we recommend that gRNA synthesis reactions be performed with our optimized one-pot reactions, either containing Klenow Fragment and RiboMAX T7 Express Enzyme Mix with 3x template oligos (either at full-scale or scaled-down), or containing the EnGen kit with 3x template oligos. We calculate that the enzymatic cost for each gRNA synthesis method, based on current vendor list-prices, to be:

(a) One-pot standard, $8.68 per gRNA from:

- Klenow Fragment (NEB, M0212L), $0.63 per gRNA ($252 / 1000 U; 2.5 U / gRNA)
- T7 RiboMAX (Promega, P1320), $8.68 per gRNA ($434 / 50 reactions)

(b) One-pot scaled-down, $2.33 per gRNA from:

- Klenow Fragment (NEB, M0212L), $0.16 per gRNA ($252 / 1000 U; 0.625 U / gRNA)
- T7 RiboMAX (Promega, P1320), $2.17 per gRNA ($434 / 200 reactions)

(c) EnGen kit, $20.80 per gRNA from:

- EnGen sgRNA Synthesis Kit (NEB, E3322S), $416 / 20 reactions

Given the flexibility of modifying components in the one-pot gRNA synthesis reactions[2], we anticipate that further increases in gRNA yield along with reductions in cost are achievable. Furthermore, sufficient yields of most gRNAs can likely be generated with one-pot incubation times shorter than 4 hours (**Sup. Fig. 15b**). Although we did not formally test scaling down the EnGen kit, it's possible that similar optimizations could be performed to comparatively reduce the cost per gRNA. Finally, most institutions have quotes from vendors that would further reduce these costs.

**Supplementary Note 4: Design considerations and recommendations for SpRYgests**

In human cells we previously observed a preference by SpRY to target sites with NRN PAMs more efficiently than those with NYN PAMs[3] (where R is A or G and Y is C or T). However, *in vitro* SpRY is essentially PAMless and can efficiently cleave sites with any PAM (**Fig. 1e**, **1f**, and **Sup. Fig. 5c**). Therefore, when designing gRNAs to perform a SpRYgest, we recommend focusing solely on the exact position at which the DSB is to be introduced. Assuming that SpRY generates a blunt DSB 3 bp upstream of the PAM, gRNAs should be designed to position the intended DSB between position 3 and 4 of the guide sequence, counting from the PAM proximal end (**Sup. Fig. 6a**). Finally, for each intended DSB two potential gRNAs can be designed, each targeting a different strand. Thus, two distinct gRNAs can be designed for each cut site, allowing the selection of the most efficient gRNA for DSB generation, and potentially avoiding rare off-target edits against closely matched sites (which can be identified and avoiding by using SpOT-check).

Once the gRNA target sites are identified, the next step is to design target-specific ~ 54 bp oligos required to generate the dsDNA template for IVT (see **Supplementary Note 1**). This workflow enables the rapid synthesis of new gRNAs and allows custom SpRYgest of any DNA sequence. Using a fixed amount of the SpRY and gRNA ribonucleoprotein (RNP) complex (with gRNAs generated as described in **Supplementary Notes 1** and

3), we were able to introduce all intended DSBs with at least >78% cleavage efficiency for all unique sites that we sought to generate DSBs (Fig. 4c). Since it is trivial to include additional SpRY RNP in the molecular reaction, we envision that the total cleavage percentage of nearly any substrate could reach completion with minor alterations to the reaction setup.

To SpRYgest 800 fmol of a supercoiled plasmid that is ~8,000 bp (approximately 4 µg of DNA), we recommend the following starting conditions that can be appropriately modified: SpRY protein at a final concentration of 1 µM and IVT gRNA (prepared without DNase treatment; see Supplementary Note 1) at a final concentration of 2 µM in Buffer 3.1 (NEB), and allowing the reaction to proceed at 37 °C for approximately 3 hours. Note, if generating gRNAs via the one-pot method, it is also possible to add 5 µL of unpurified and unquantified gRNA IVT reaction directly into the SpRYgest reaction mixture (which generally will lead to the inclusion of a large excess of the gRNA). If incomplete SpRYgestion is observed, we recommend performing a clean-up of the gRNA preparation and proceeding with quantification to ensure that adequate gRNA is being generated. Finally, the timing of gRNA synthesis via the one-pot method can be varied and reduced to as little as 1 hour for most gRNAs (Sup. Fig. 15b and 15c).

**Supplementary Note 5: Annotation of substrate off-targets via SpOT-check**

To identify potential gRNAs with potential closely matched off-target sites in prospective DNA substrates, we developed software called SpOT-check (SpRYgest off-target checker), which is freely available online at www.kleinstiverlab.org/spotcheck. Beyond enumerating the number and sequences of closely matched sites in the substrate, SpOT-check can also be used to design gRNA sequences to create a cut at a designated position (for both top and bottom strand, with oligos needed for IVT as output) in a provided plasmid sequence. Alternatively, pre-designed guide sequences can be used as the input. For each gRNA sequence, SpOT-check reports any potential off-target sites up to off-by-6 mismatches, permitting users to prioritize gRNAs targeting the strand with fewer potential off-targets. SpOT-check "batch mode" will also be available for download from GitHub, allowing users to input multiple guides and plasmid sequences at once in CSV format as well as specifying any maximum number of mismatches for which to report off-targets. Using SpOT-check, we computed the off-target profiles of all gRNAs used in this study (Sup. Table 5).

#### Supplementary Figures:

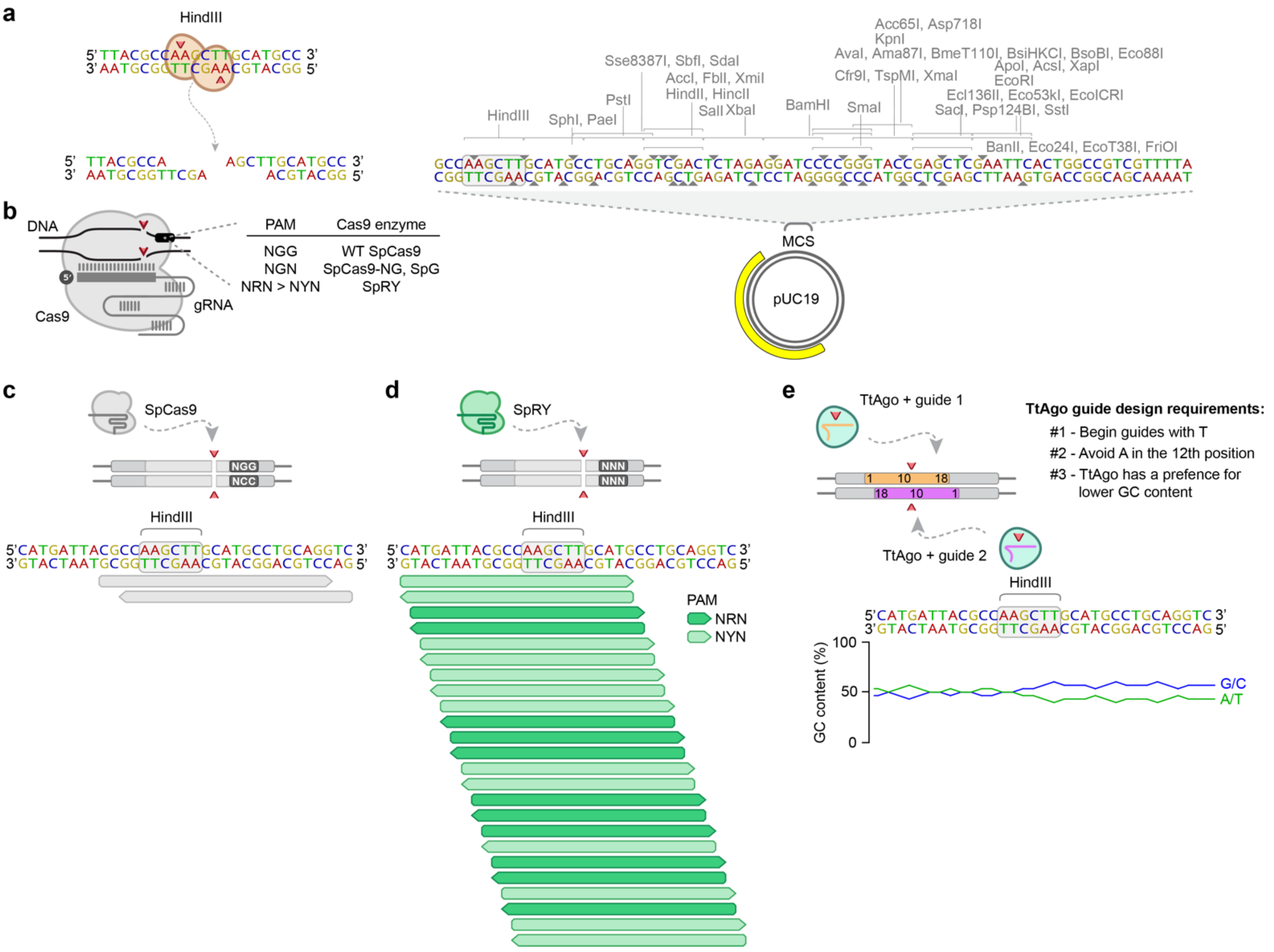


**Supplementary Figure 1: Comparison of DNA-targeting endonucleases.** (**a**) Illustration of target recognition and cleavage by HindIII (left panel) and of the multiple cloning site (MCS) in pUC19 with any single cutter restriction enzyme shown, including HindIII (right panel). (**b**) Schematic of SpCas9 complexed with a gRNA and bound to a target DNA sequence bearing various PAMs, depending on the SpCas9 variant. gRNA, guide RNA; PAM, protospacer-adjacent motif. (**c**) The PAM requirement of wild-type (WT) SpCas9 restricts target site design and DNA cleavage to sites harboring NGG PAMs, limiting precise DNA break positioning in a DNA molecule. (**d**) In contrast to WT SpCas9, the near-PAMless SpRY variant might offer unconstrained targeting for *in vitro* DNA digests. For panels c and d, target sites (20 nt spacer with 3 nt PAM) are shown that target the top and bottom strand (with right and left facing points, respectively). (**e**) Schematic of guide design requirements to potentially generate a double strand break with TtAgo.

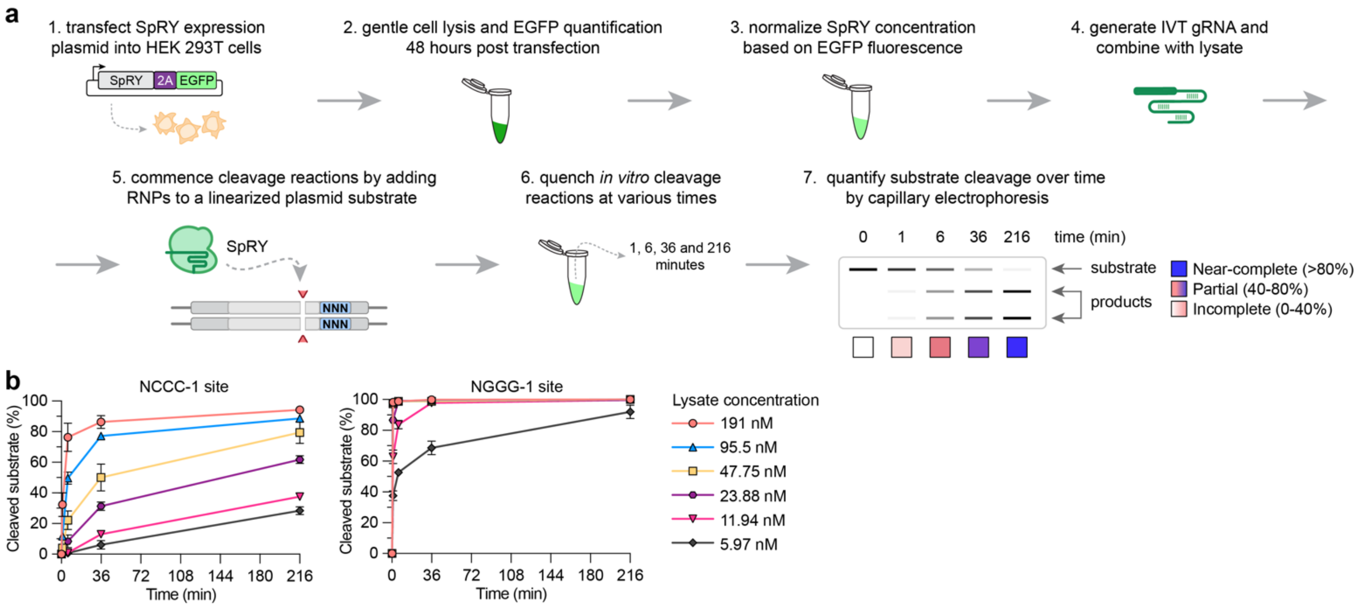


**Supplementary Figure 2: Generation of SpRY protein from human cell lysate for SpRYgest cleavage reactions.** (**a**) Schematic of SpRY production and SpRYgest workflow. SpRY protein is expressed in human cells and harvested by gentle lysis, with SpRY concentrations normalized by EGFP fluorescence versus a fluorescein standard curve. The SpRY protein from the lysate is complexed with a gRNA (produced by *in vitro* transcription or via chemical synthesis), and the resulting RNPs are combined with a linearized plasmid template to initiate *in vitro* cleavage reactions. Aliquots from the reaction are extracted at various timepoints, terminated, and the cleavage of DNA substrates over time is analyzed by capillary electrophoresis. (**b**) Titration of normalized lysate to vary the amount of SpRY protein as input into *in vitro* cleavage reactions, performed utilizing gRNAs targeted to NCCC-1 or NGGG-1 sites (left and right panels, respectively). Unless otherwise noted, all other small-scale SpRYgests using linearized plasmid substrates were performed using 180 nM fluorescein-normalized SpRY lysate. The cleavage of DNA substrates was analyzed by capillary electrophoresis; mean and s.e.m. shown for n = 3.

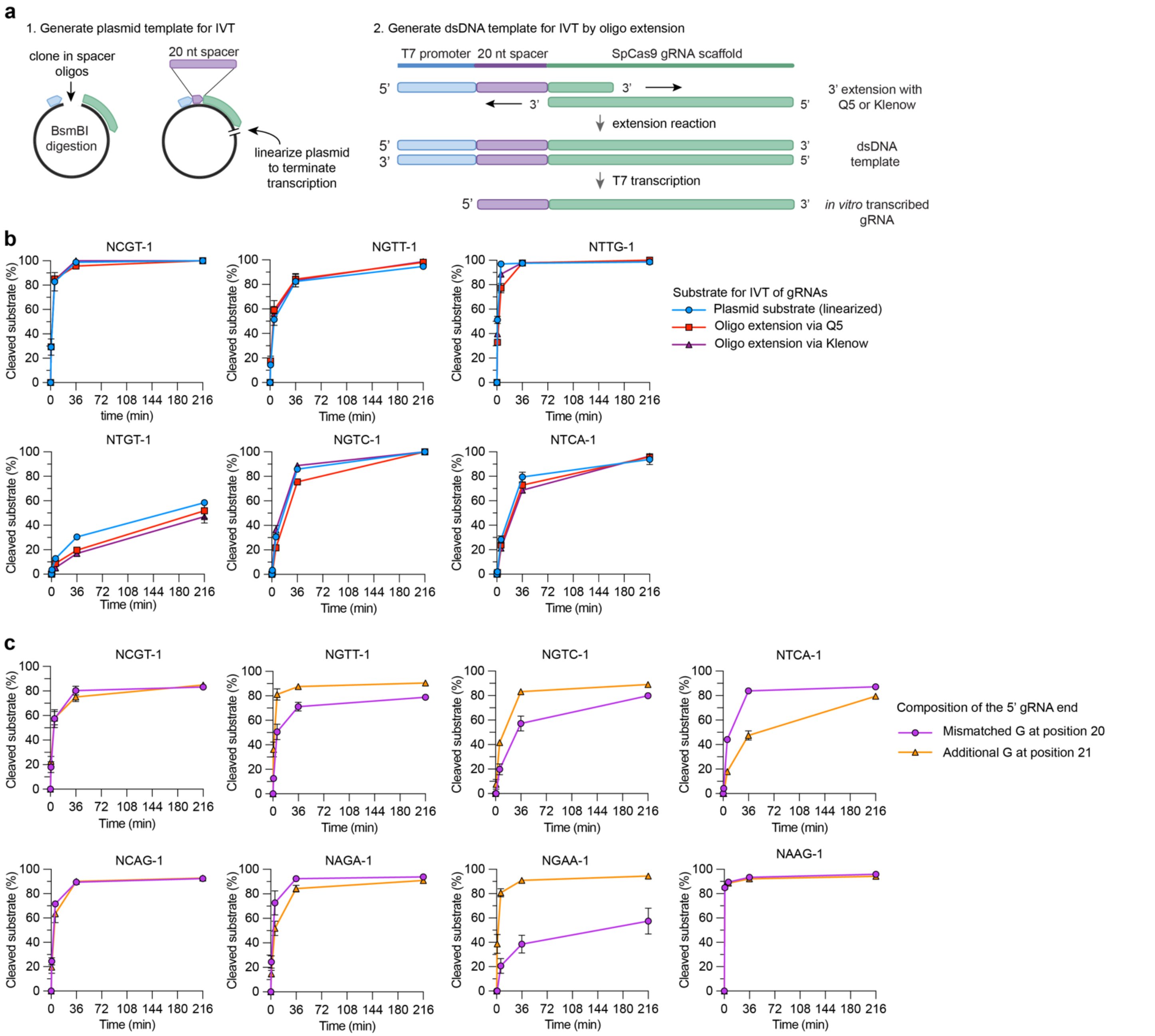


**Supplementary Figure 3: gRNA production via *in vitro* transcription.** (**a**) Illustration of three different methods to produce gRNAs via *in vitro* transcription (IVT), using templates generated either by (1) cloning a dsDNA plasmid template with a T7 promoter, the spacer sequence, and gRNA scaffold (left panel), or (2) extension of overlapping ssDNA oligonucleotides using either Q5 polymerase or Klenow (right panel). (**b**) Comparison of the effectiveness of SpRYgests when using gRNAs produced via IVT from the three different DNA substrates shown in **panel a** (run-off IVT from a cloned plasmid, or from a dsDNA fragment generated by oligonucleotide extension with Q5 or Klenow). (**c**) Comparison of the effectiveness of SpRYgests when using gRNAs transcribed with 20 or 21 nt spacers (containing a matched or mismatched 5'G to improve transcription from the T7 promoter). For panels b and c, the cleavage of DNA substrates was analyzed by capillary electrophoresis; mean and s.e.m. shown for n = 3.

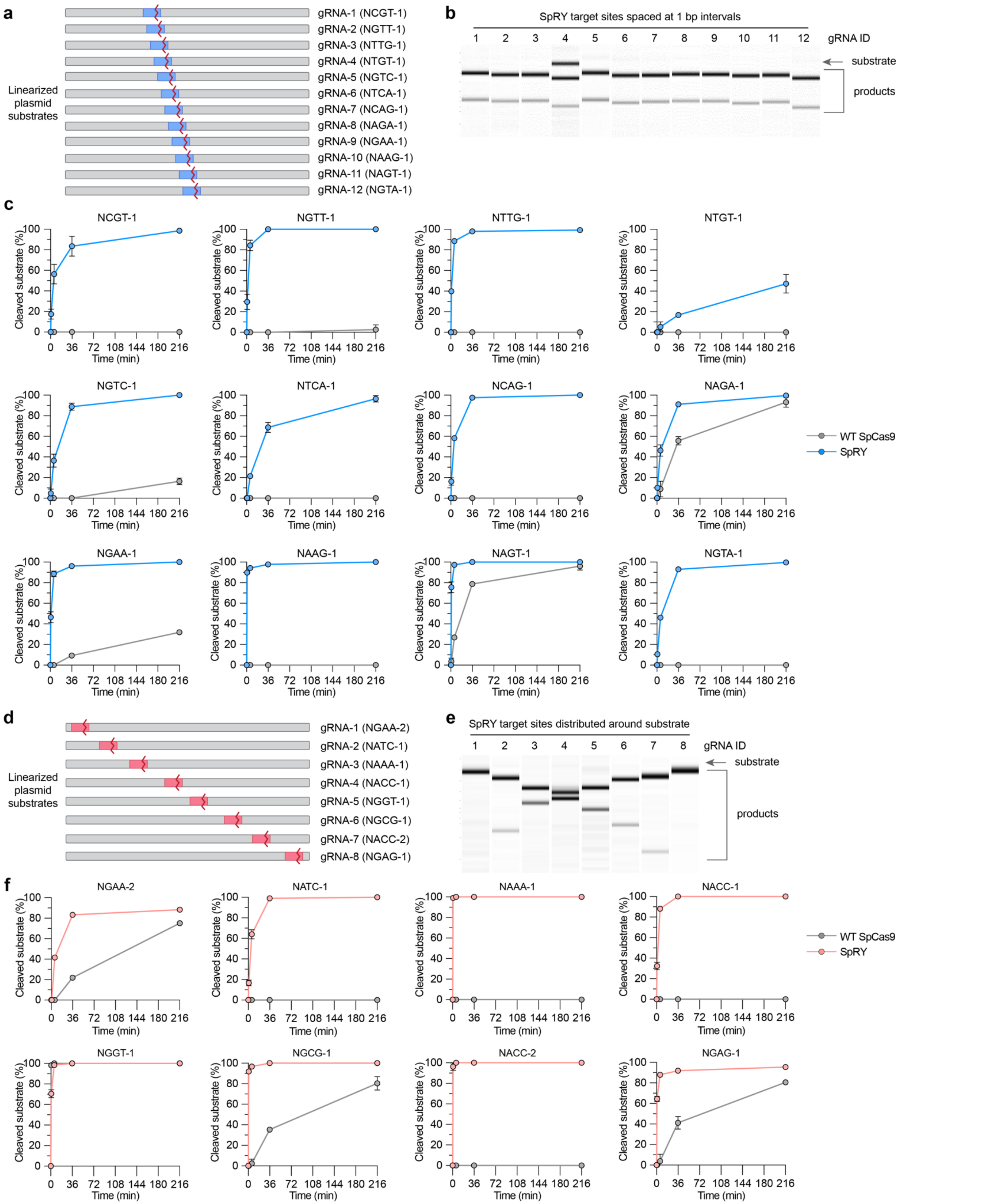


**Supplementary Figure 4: Initial SpRYgest cleavage experiments on 20 sites.** (**a**) Schematic of the 12 gRNAs in a specific region of the plasmid that shift by 1 bp intervals. (**b**) Visualization of the cleavage products of

SpRYgest reactions at the final timepoint (216 minutes) when using the 12 1-nt interval gRNAs and lysates containing SpRY, run on a QIAxcel Fast Analysis Kit (Qiagen). (**c**) Plots of substrate cleavage over time with the 12 1bp-spaced gRNAs in the presence of lysates containing WT SpCas9 or SpRY. (**d**) Schematic of the 8 gRNAs distributed at various positions in the substrate. (**e**) Visualization of the cleavage products of SpRYgest reactions at the final timepoint (216 minutes) when using the 8 distributed gRNAs and lysates containing SpRY, run on a QIAxcel Fast Analysis Kit. (**f**) Plots of substrate cleavage over time with the 8 distributed gRNAs in the presence of lysates containing WT SpCas9 or SpRY. For **panels c** and **f**, cleavage of DNA substrates was analyzed by capillary electrophoresis; mean and s.e.m. shown for n = 3.

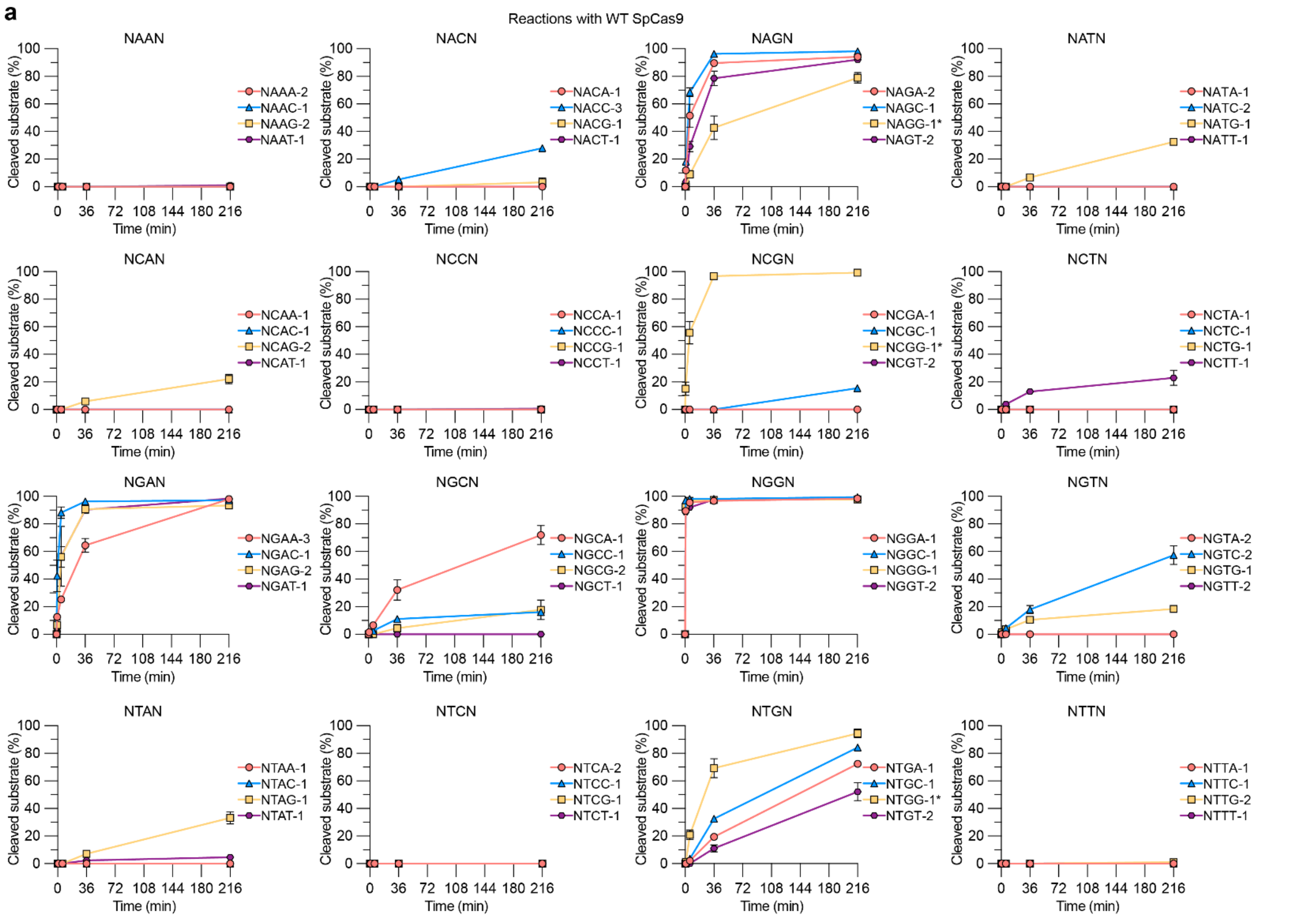


***Figure 5a*** *(legend follows)*

**b**

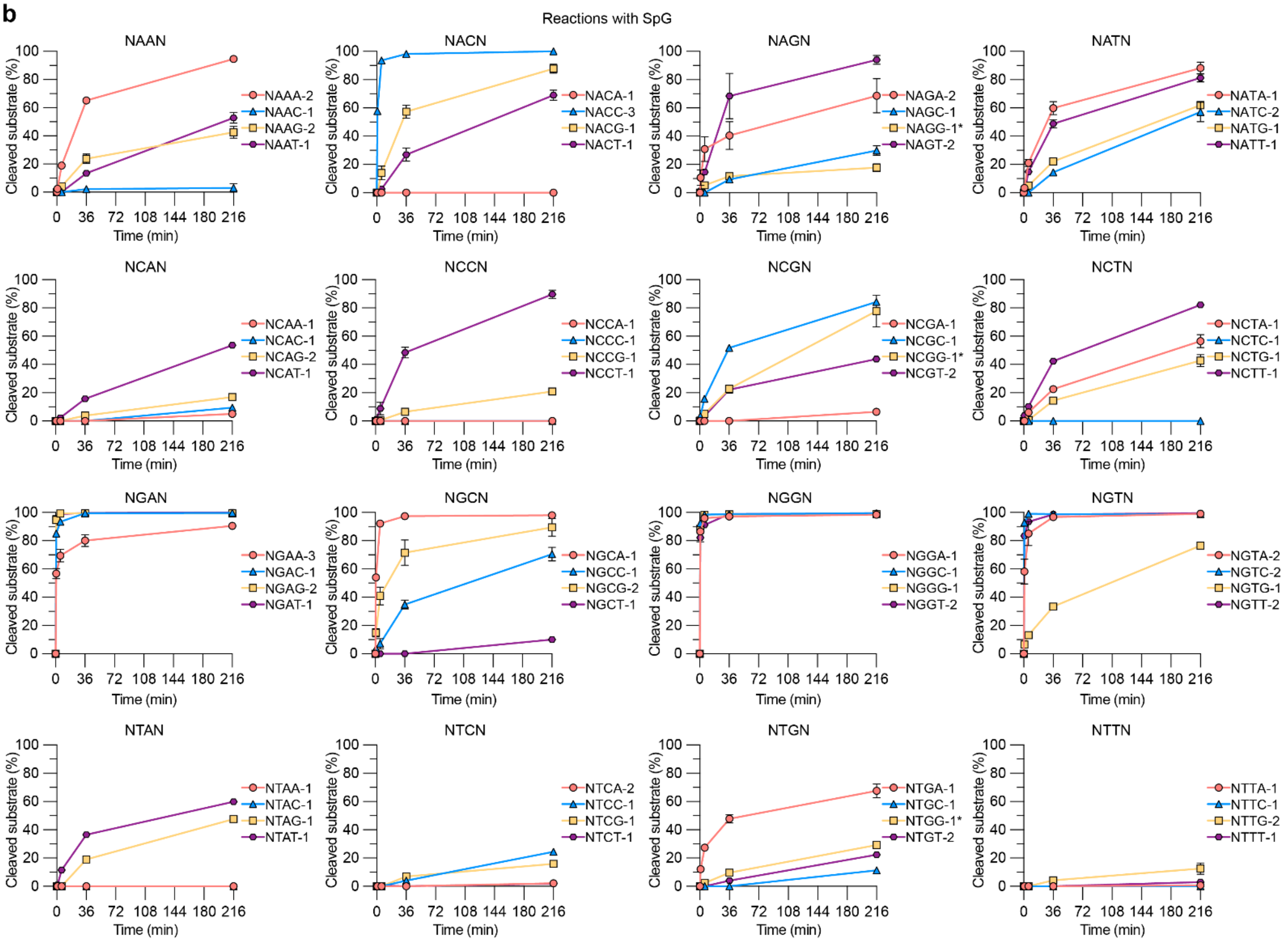


***Figure 5b*** *(legend follows)*

**c**

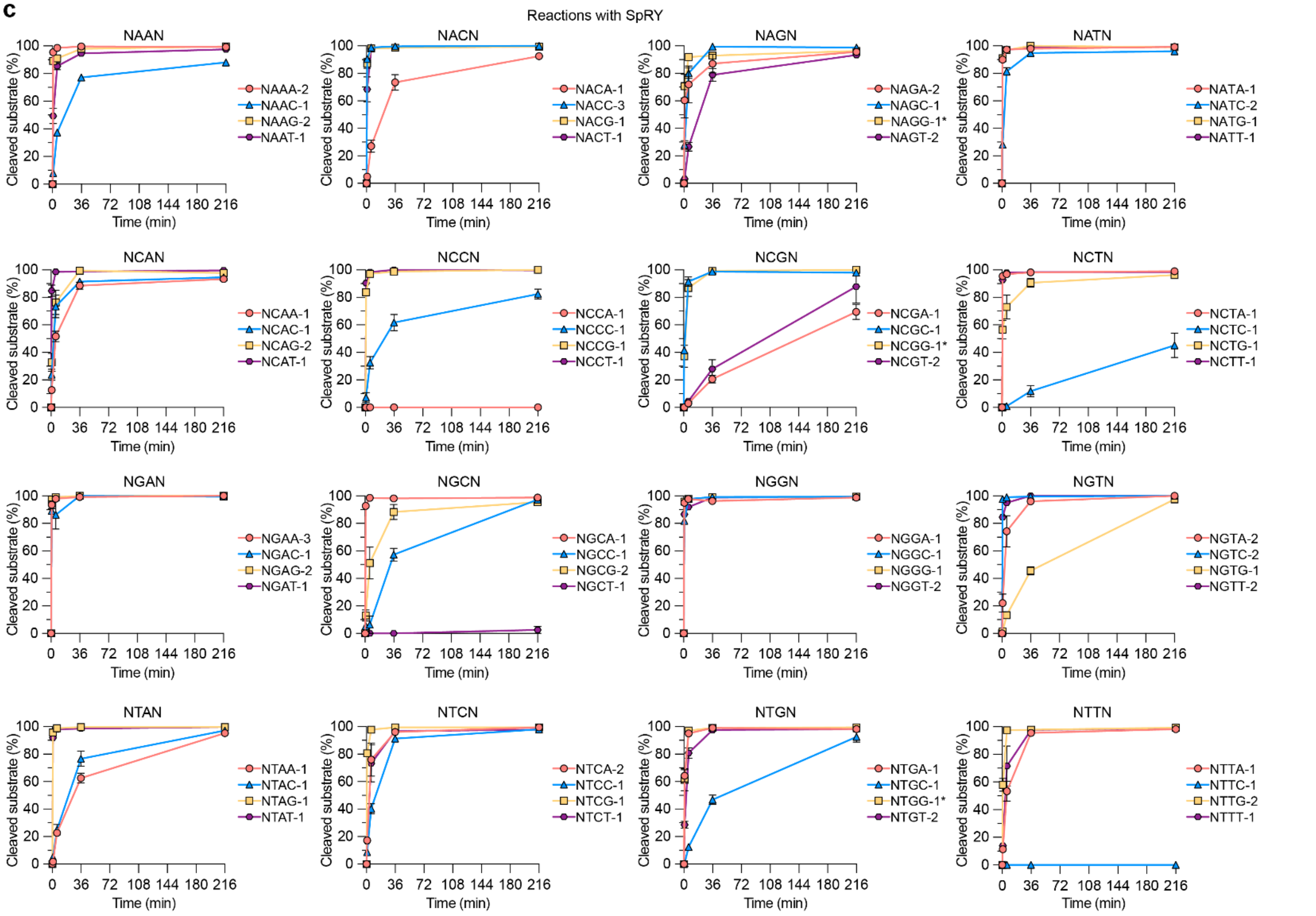


**Supplementary Figure 5: Cleavage activity of WT, SpG, and SpRY across 64 sites.** (**a-c**) Time-course *in vitro* cleavage reactions were performed using wild-type (WT) SpCas9, SpG, or SpRY[3] protein generated from human cell lysates (**panels a-c**, respectively). Lysates were separately combined with 64 different gRNAs targeted to sites that sample each possible identity in the 2nd, 3rd, and 4th positions of an NNNN PAM, as well as a linearized plasmid substrate. Cleavage of DNA substrates was analyzed by capillary electrophoresis; mean and s.e.m. shown for n = 3.

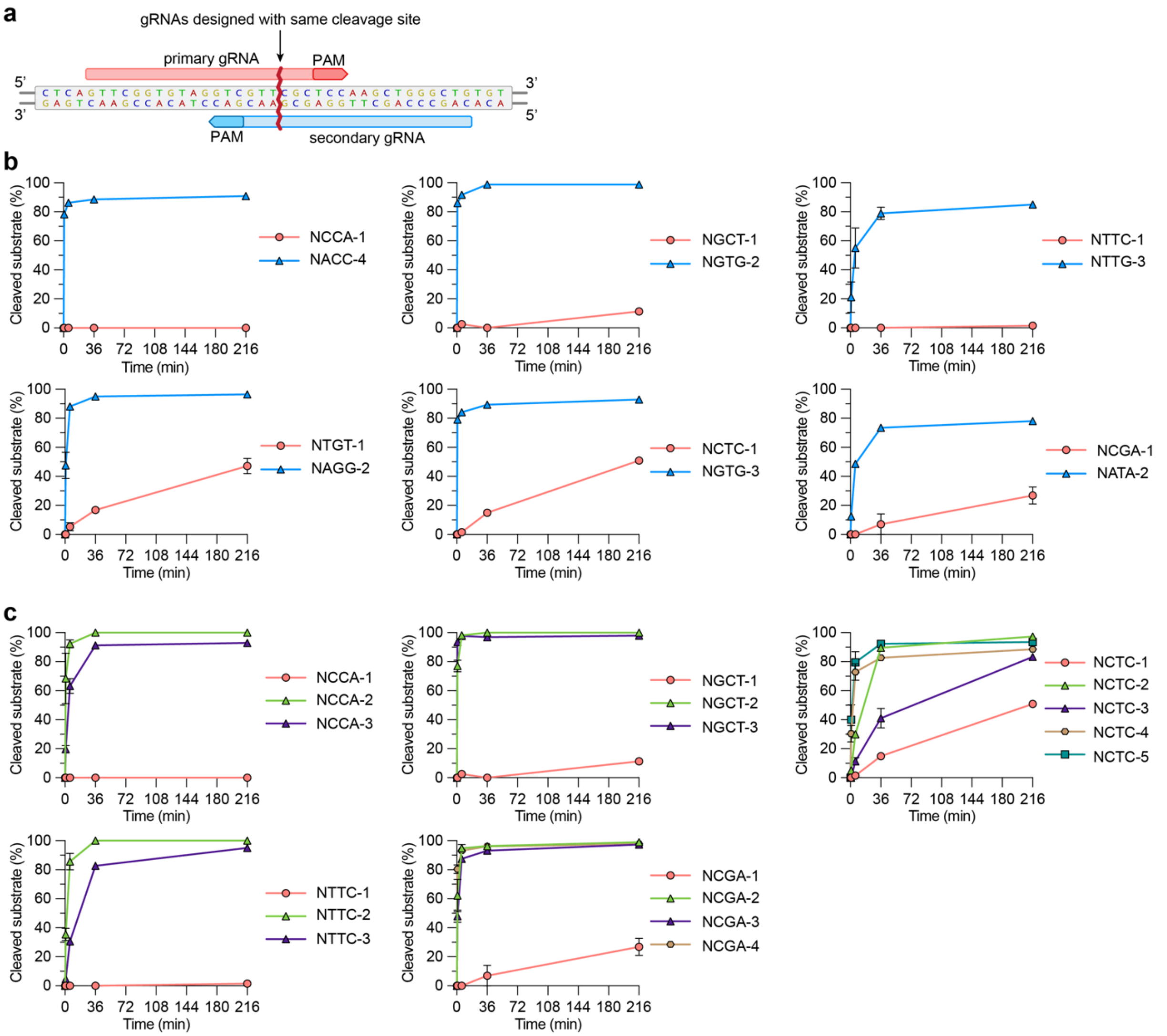


**Supplementary Figure 6: Reconciliation of failed SpRYgest sites.** (**a**) Schematic of utilizing a secondary gRNA to generate the same DSB as the primary gRNA by targeting the opposite strand, based on SpRY generating a blunt DSB 3 nt from the PAM. (**b**) Comparison of the cleavage efficiencies of primary gRNAs that exhibited partial or incomplete cleavage, and newly designed secondary gRNAs targeted to the opposite strand. (**c**) Cleavage efficiencies of supplementary gRNAs designed to target additional sites with the same PAM as the primary gRNAs that exhibited partial or incomplete cleavage. For **panels b** and **c**, an asterisk (*) indicates the primary gRNA; cleavage of DNA substrates was analyzed by capillary electrophoresis; mean and s.e.m. shown for n = 3.

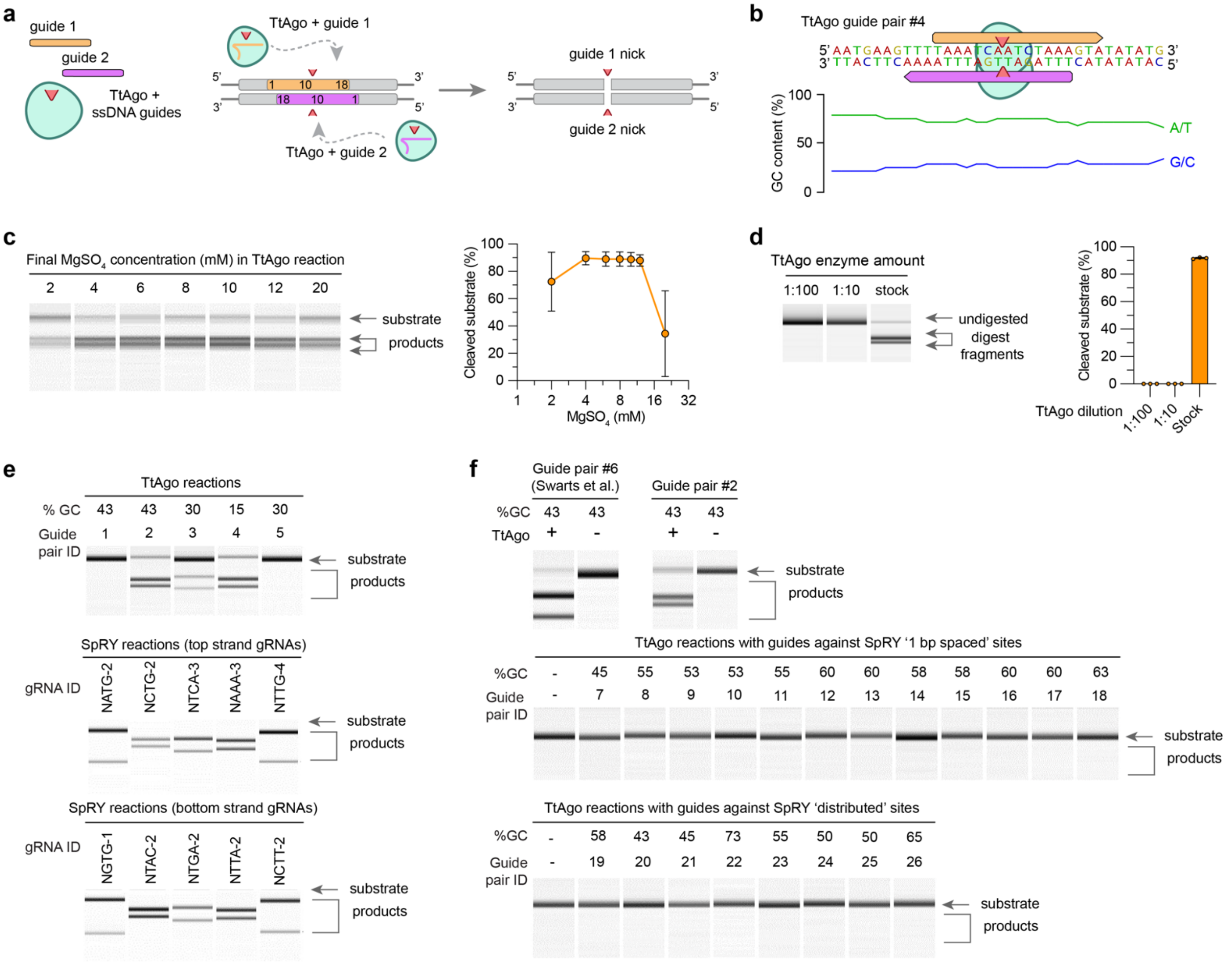


**Supplementary Figure 7: Substrate cleavage reactions with TtAgo.** (**a**) Schematic of TtAgo generating a DSB when programmed with two separate 5'P-ssDNA guides. (**b**) Schematic of TtAgo guide design for a site that was compatible with the design requirements (TtAgo guide pair #4; see **Fig. 2a** and **Supplementary Table 4**). (**c**) Titration of $MgSO_4$ concentration in TtAgo cleavage reactions using a linearized substrate with a pair of 5'P-ssDNA guides to generate a DSB (guide pair #2; see **Supplementary Table 4**). (**d**) Titration of TtAgo enzyme concentration in cleavage reactions performed on a linearized substrate with a pair of 5'P-ssDNA guides (guide pair #2). For **panels c** and **d**, the left panels are representative traces from one replicate generated using QIAxcel ScreenGel Software v1.5 (Qiagen), and the right panels are graphs showing data from 3 independent replicates. (**e**) Capillary electrophoresis traces comparing final timepoint aliquots of substrate cleavage reactions with TtAgo and SpRY on 5 sites (at 60 and 216 minutes, respectively), as shown in **Fig. 2a**. These 5 sites follow the TtAgo guide design requirements (see **Supplementary Note 2**). (**f**) Capillary electrophoresis traces showing final timepoint aliquots of substrate cleavage reactions with TtAgo and pairs of 5'P-ssDNA guides (analyzed after 60 minute reactions). Two guide pairs (with pair #6 from Swarts et al.[1] and #2 from **Fig. 2a**) were previously shown to result in near-complete cleavage (top panel), and 20 new sites that do not follow the TtAgo guide design requirements (analogous to the SpRYgest sites from **Figs. 1c** and **1d** in the middle and bottom panels, respectively). For **panels c-f**, cleavage of DNA substrates was analyzed by capillary electrophoresis; with mean and s.e.m. shown for n = 3 in graphs or n = 1 in traces.

**a**

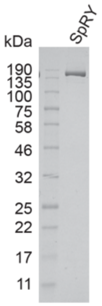


**Supplementary Figure 8: Purification of SpRY.** (**a**) Representative gel image of purified SpRY protein run alongside a protein standard (Broad Range; New England Biolabs P7712). Novex 10-20% Tris-glycine gel post stained with SimplyBlue Safe Stain (both purchased from ThermoFisher).

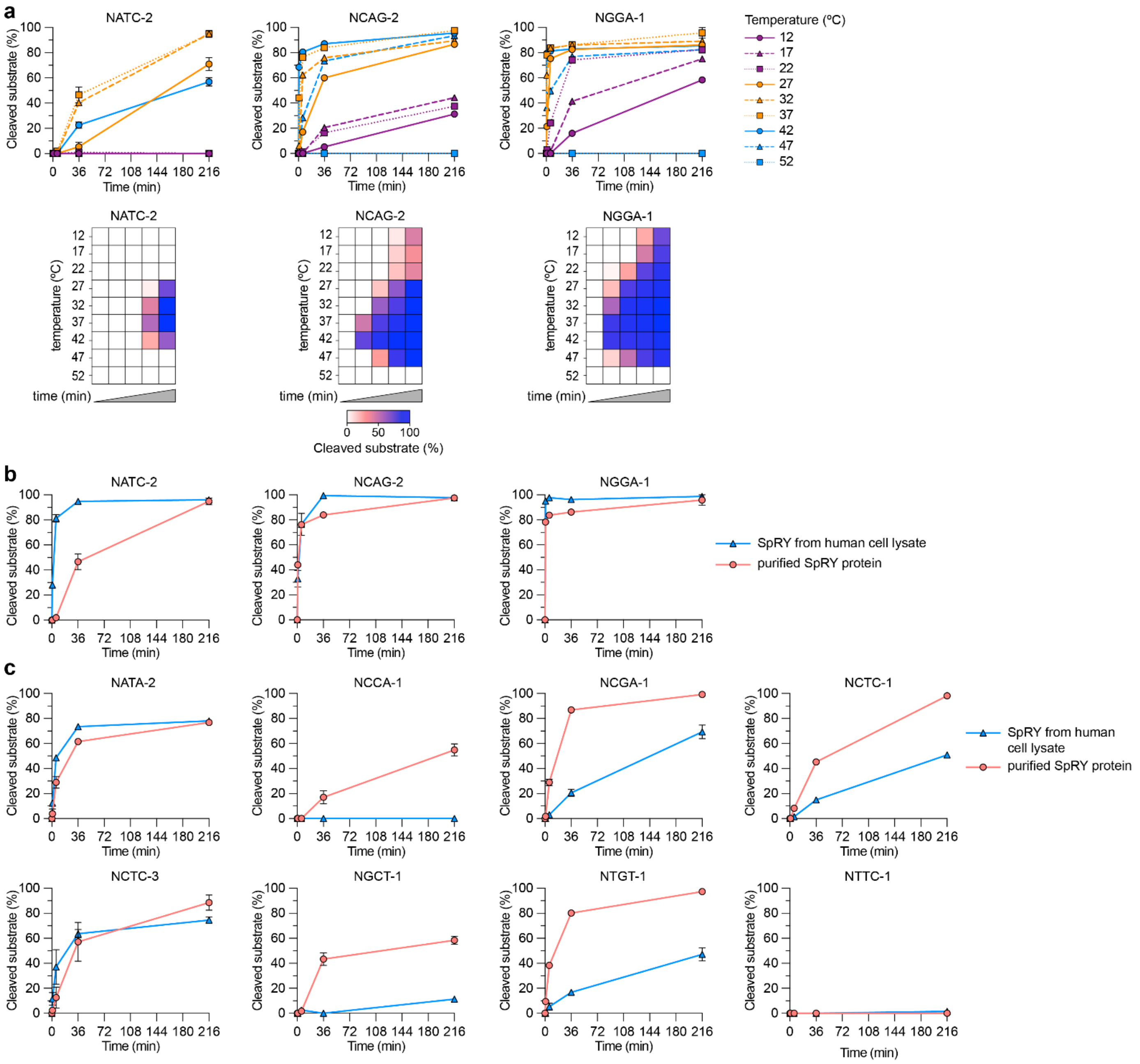


**Supplementary Figure 9: Assessment of SpRYgest parameters.** (**a**) Time-course SpRYgests performed using SpRY expressed from human cell lysates were performed across nine different temperatures, plotted as scatterplots or heatmaps (top and bottom panels, respectively). (**c**,**d**) Plots illustrating the rates of substrate cleavage when using SpRY from human cell lysates or purified SpRY protein, for gRNAs whose reactions with SpRY lysates either went to completion (**panel b**) or exhibited partial or incomplete substrate cleavage (**panel c**). Reactions were performed at 37ºC. For **panels a**-**c**, cleavage of DNA substrates was analyzed by capillary electrophoresis; mean, and in some cases s.e.m., shown for n = 3.

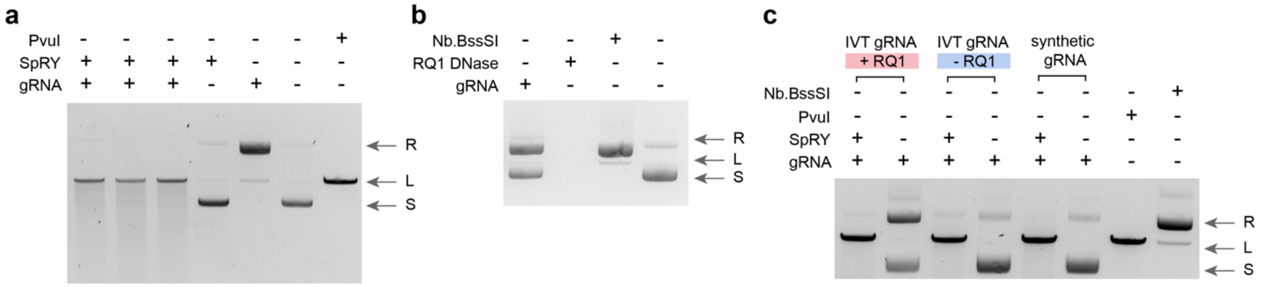


**Supplementary Figure 10: DNase contamination during gRNA preparation can cause non-specific degradation of substrate.** (**a**) Agarose gel illustrating degradation of plasmid substrates, resulting from a probable carry-forward of the RQ1 DNase present in the gRNA *in vitro* transcription protocol. (**b**) Agarose gel illustrating plasmid conformations after incubation with the a gRNA only control, concentrated RQ1 DNase (Promega), or Nb.BssSI nickase (New England Biolabs). (**c**) Agarose gel comparing plasmid conformations after treatment with RNPs formed with *in vitro* transcribed gRNAs generated with and without the RQ1 DNase treatment, and commercial synthetic gRNAs. For all panels, plasmid conformations are: R, relaxed; L, linear; S, supercoiled. Nb.BssSI is a nickase; all agarose gels were stained with ethidium bromide.

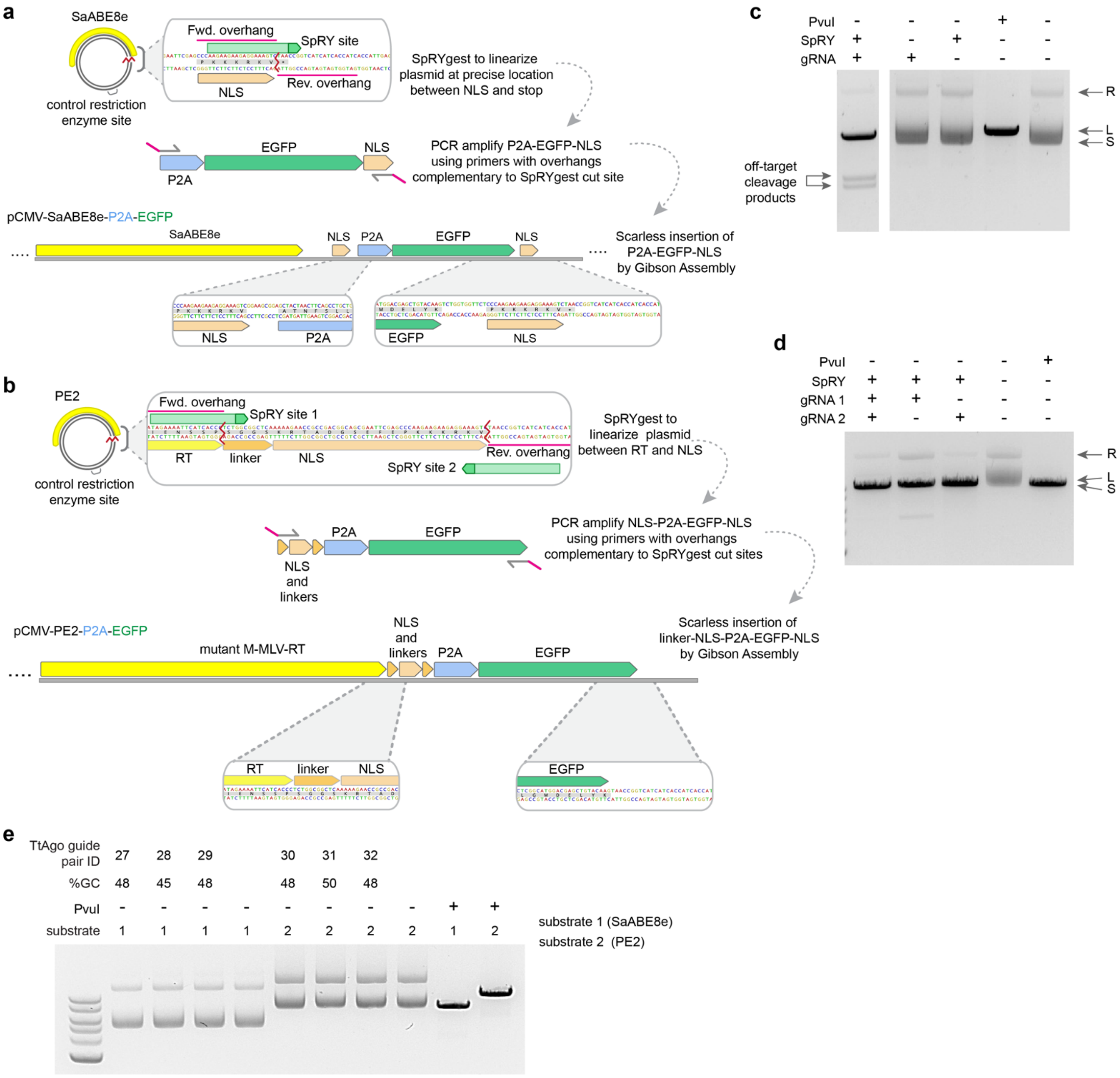


**Supplementary Figure 11: Cloning reactions to generate large sequence insertions via SpRYgest.** (**a**,**b**) Schematics of the SpRYgest cloning strategies to add P2A-EGFP sequences to SaCas9-ABE8e and PE2 plasmids via single and double SpRYgests, in **panels a** and **b**, respectively. (**c**,**d**) Agarose gel images for the SpRYgest reactions of SaCas9-ABE8e and PE2 plasmids in **panels c** and **d**, respectively. (**e**) Agarose gel image of TtAgo reactions with pairs of 5'P-ssDNA guides targeted to the same SpRYgest sites in **panels a** and **b**. For **panels c**-**e** plasmid conformations are: R, relaxed; L, linear; S, supercoiled; gels were stained with ethidium bromide; the ladder in **panel e** is the 1 kb Plus DNA Ladder (New England Biolabs).

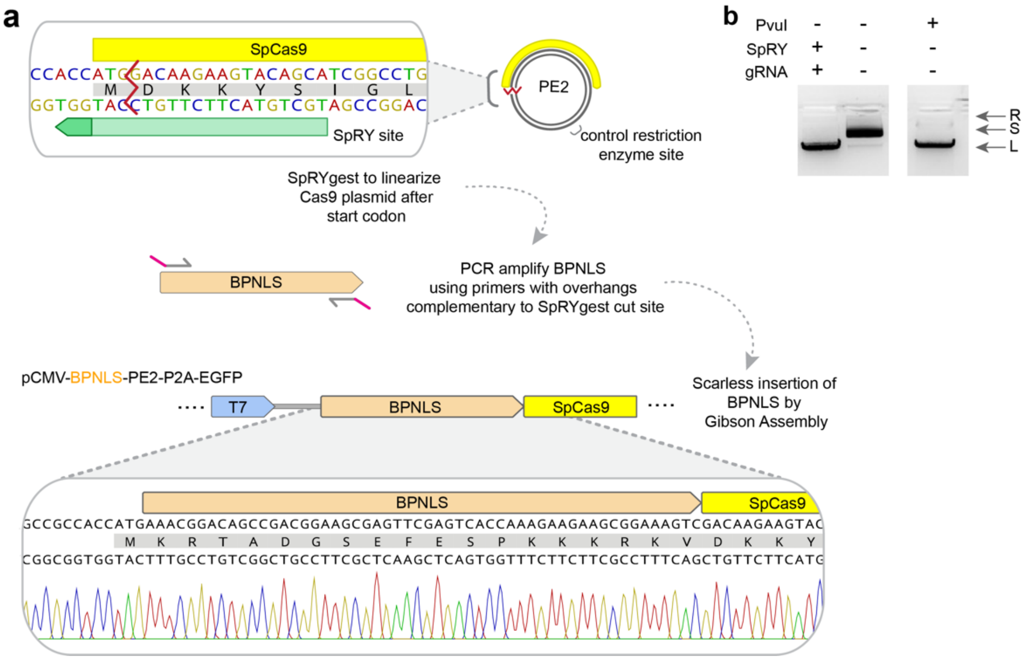


**Supplementary Figure 12: Short sequence insertions via SpRYgest.** (**a**) Schematic of the SpRYgest strategy to add a short sequence (a BPNLS) to the N-terminus SpCas9 in a human expression plasmid. The section of a representative Sanger sequencing reaction illustrates the cloning junction from a correct clone. (**b**) Agarose gel image for the SpRYgest reaction in panels a. Plasmid conformations are: R, relaxed; L, linear; S, supercoiled; gel stained with ethidium bromide.

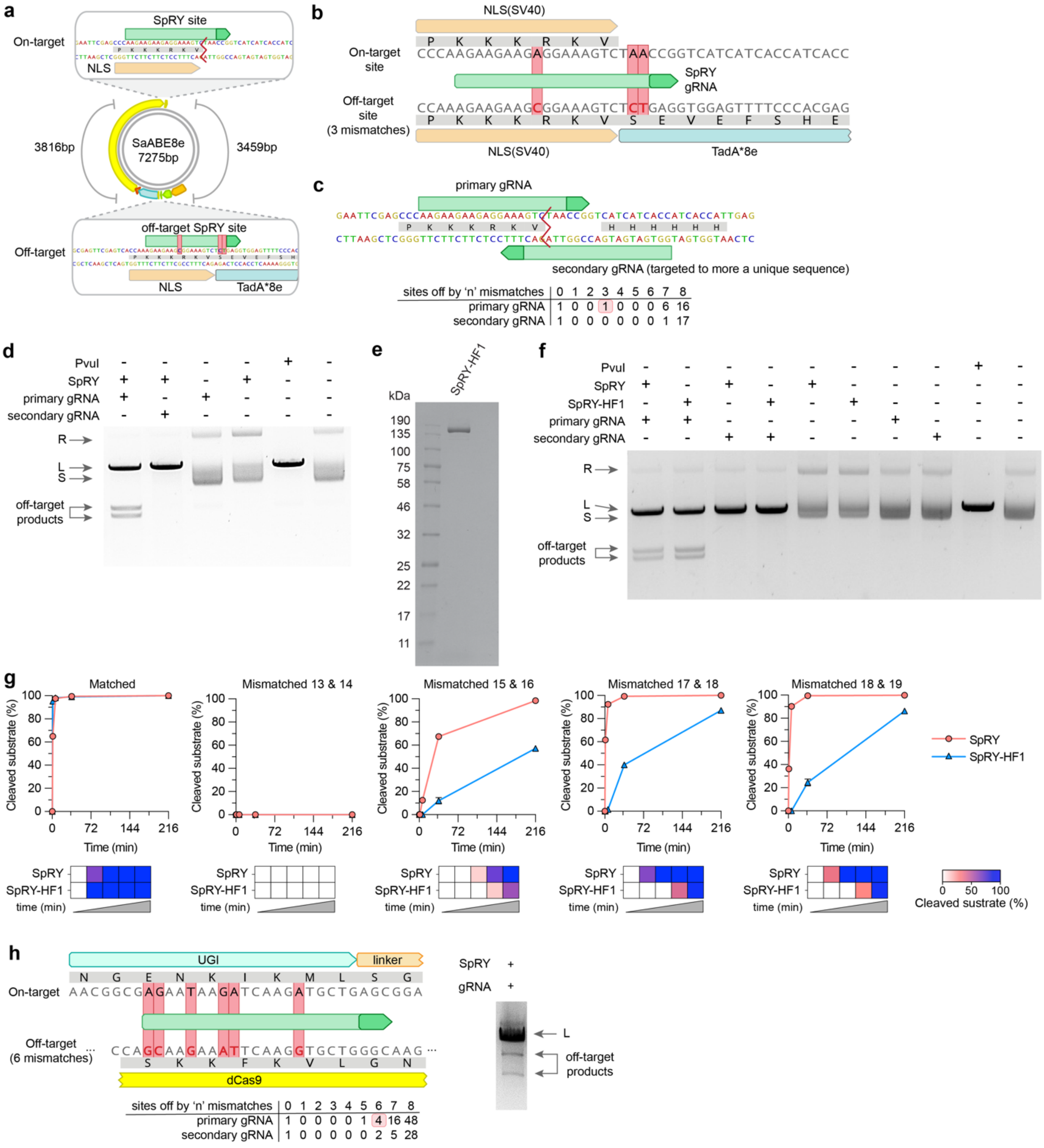

**Supplementary Figure 13: Observation and mitigation of off-target cleavage in SpRYgest reactions.** (**a**) Schematic of the SaCas9-ABE8e plasmid used in **Sup. Fig. 11a**, with the intended on-target site and an off-target site containing 3 mismatched bases highlighted. (**b**) Sequence alignment illustrating the positions of mismatches between the SaCas9-ABE8e on-target and off-target sites. (**c**) Schematic showing the strategy of targeting the opposite strand with a secondary gRNA in the SpRYgest of the SaCas9-ABE8e plasmid. The bottom panel table shows the number of putative closely matches sites in the target plasmid, generated using SpOT-check. The red shading highlights the off-by-3 mismatch off-target that led to cleavage by SpRY. (**d**) Agarose gel image showing the mitigation of off-target SpRYgest cleavage of the SaCas9-ABE8e plasmid by targeting the opposite strand with a secondary gRNA. (**e**) Representative gel image of purified SpRY-HF1[3]

protein run alongside a protein standard (Broad Range; New England Biolabs P7712). Novex 10-20% Tris-glycine gel post stained with SimplyBlue Safe Stain (both purchased from ThermoFisher). (**f**) Agarose gel cleavage reactions with SpRY and SpRY-HF1 when using the primary and secondary gRNAs for the SaCas9-ABE8e plasmid digest. (**g**) Comparison of the cleavage efficiency of purified SpRY and SpRY-HF1 when using gRNAs that are either perfectly matched or that harbor pairs of mismatches to the target spacer sequence. Cleavage of DNA substrates was analyzed by capillary electrophoresis; mean, and in some cases s.e.m., shown for n = 3. (**h**) Sequence alignment of on- and off-target sites for an additional gRNA for which low level off-target cleavage was observed in SpRYgest of site NGCG-4. The bottom panel table shows the number of putative closely matches sites in the target plasmid, generated using SpOT-check. The red shading highlights the off-by-6 mismatch off-target that led to cleavage by SpRY. For **panels d**, **f**, and **h**, plasmid conformations are: R, relaxed; L, linear; S, supercoiled; gels stained with ethidium bromide

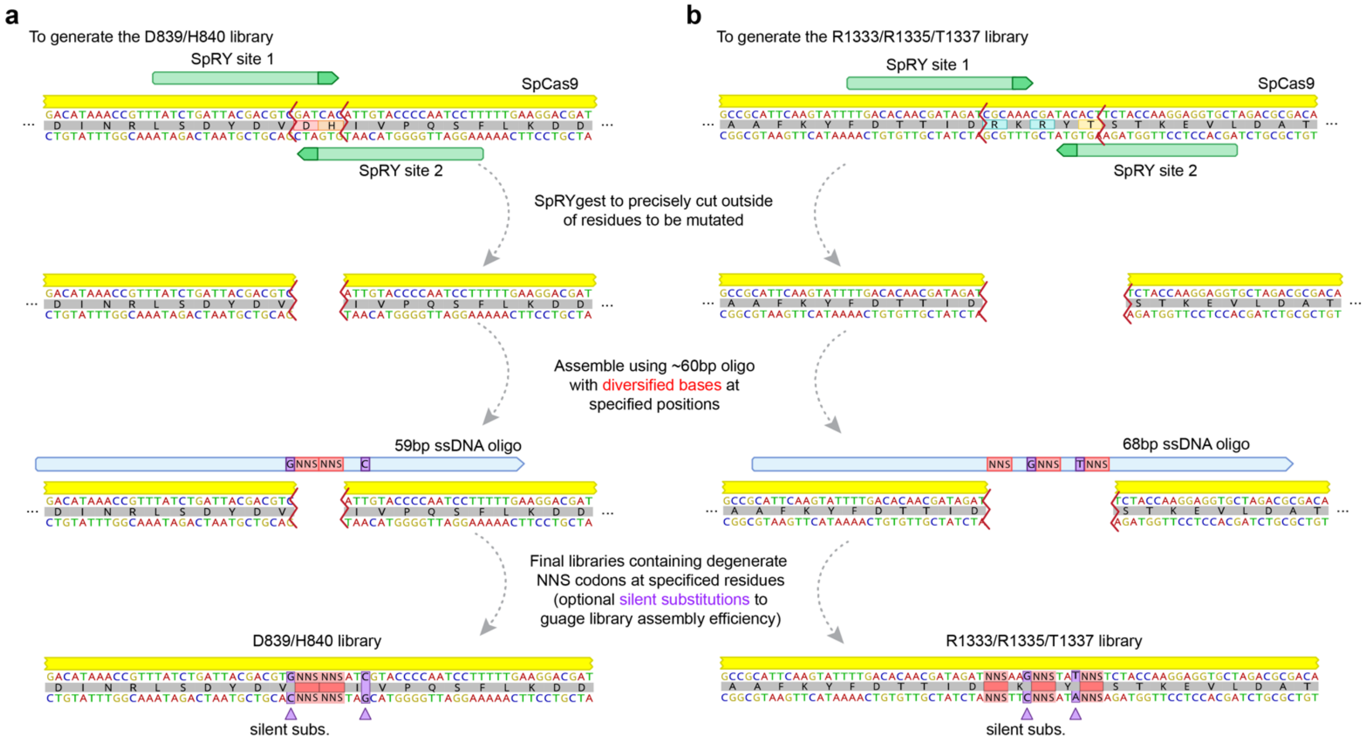


**Supplementary Figure 14: Molecular cloning of saturation mutagenesis libraries within regions of SpCas9 via SpRYgest.** (**a**,**b**) Schematics of SpRYgest strategies to generate saturation mutagenesis libraries of residues of SpCas9 important for its catalytic activity (within the HNH domain; **panel a**) and its NGG PAM preference (within the PAM-interacting domain; **panel b**). Silent substitutions were included in the library oligo to enable assessment of library construction efficiency (marked in orange boxes).

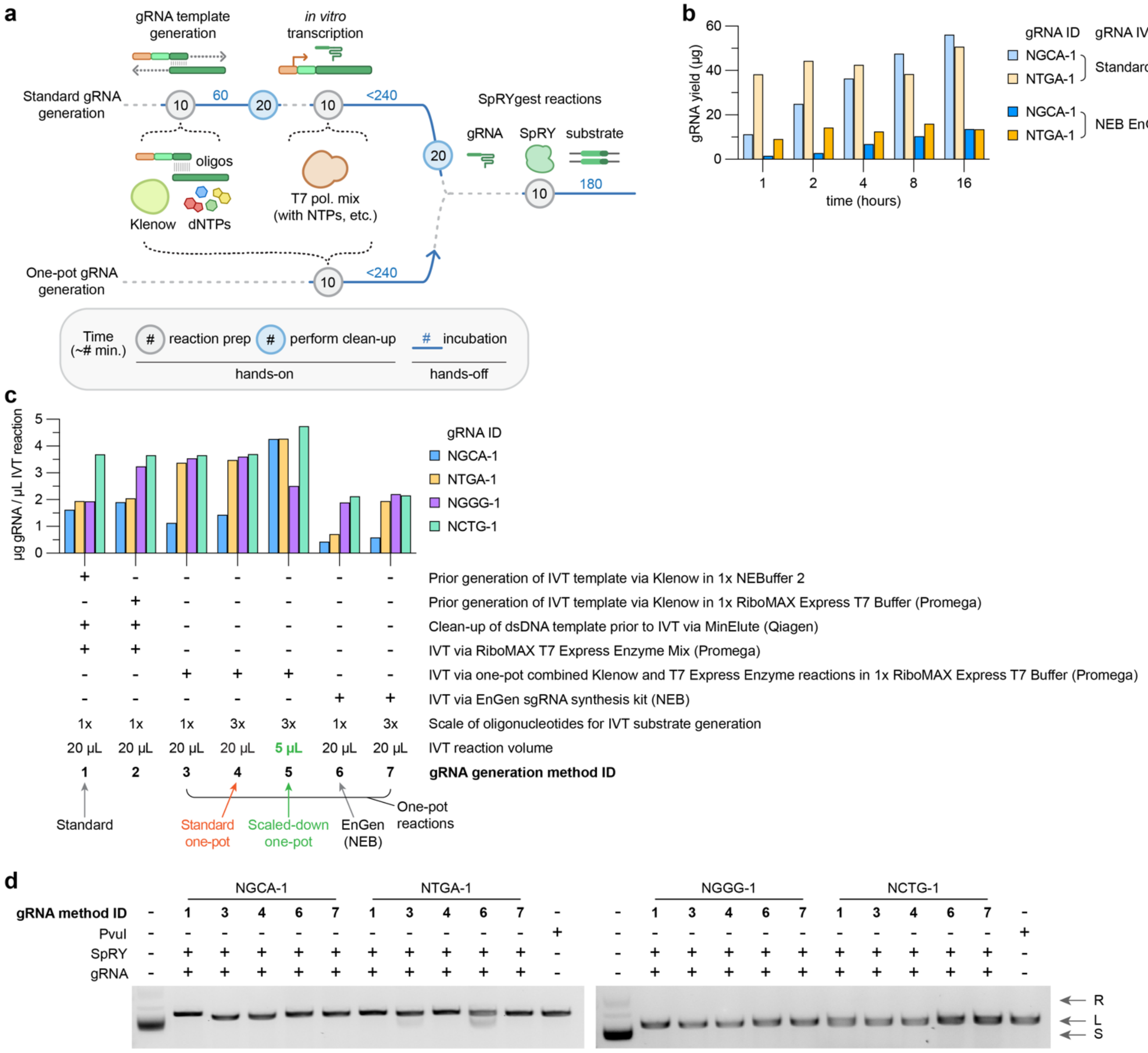


**Supplementary Figure 15: Optimization of a one-pot gRNA IVT reaction.** (**a**) Schematic illustrating the hands-on and incubation times of our previously established Standard gRNA synthesis method (Klenow + T7 RiboMAX; see panel c and Supplementary Note 1) versus streamlined 'one-pot' reactions. Approximate times in minutes are shown; lines not drawn to scale. (**b**) Comparison of *in vitro* transcribed gRNA yield for our previously established Standard gRNA synthesis method and a commercially available one-pot kit (EnGen sgRNA Synthesis Kit; NEB, # E3322S). (**c**) Comparison of *in vitro* transcribed gRNA yield generated via different methods with 4 hour reaction incubations (see also Supplementary Note 3). The gRNA yields are corrected for the µL volumes of each transcription reaction. For panels b and c, the IVT products were cleaned up using paramagnetic beads prior to quantification. (**d**) Agarose gel of SpRYgest reactions performed with gRNAs taken directly from gRNA synthesis reactions without conducting a clean-up step. The gRNA method IDs refer to the method IDs from panel c. The quantities of gRNAs NGCA-1 and NTGA-1 used in SpRYgest reactions were normalized to final concentration of 2 µM in the 50 µL SpRYgest reaction based on concentrations determined in duplicate reactions that were cleaned up and quantified (plotted in **panel c**). For SpRYgest reactions performed with gRNAs NGGG-1 and NCTG-1, 5 µL of the uncleaned and unquantified gRNAs from the one-pot reactions were used directly in the SpRYgest reaction. Plasmid conformations are: R, relaxed; L, linear; S, supercoiled. Agarose gels were stained with ethidium bromide.

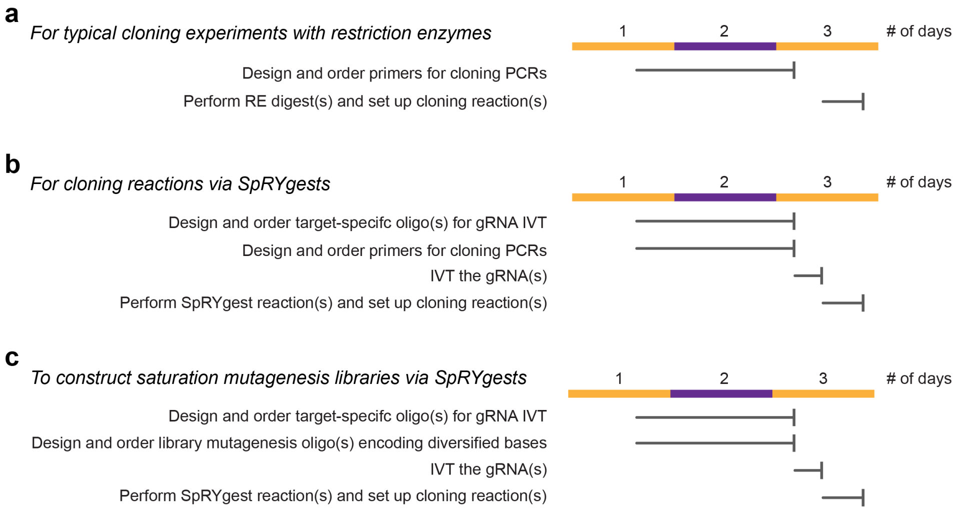


**Supplementary Figure 16: Timelines for molecular cloning reactions.** (**a**-**c**) Gant charts and workflow details for performing standard cloning reactions with restriction enzyme digests (**panel a**), cloning reactions via SpRYgests (**panel b**), and for generating saturation mutagenesis library via SpRYgests (**panel c**). Note that the oligonucleotide ordering times and most of the other process steps also apply to restriction enzyme cloning. The timing of oligo ordering can be expedited if local DNA synthesis facilities or next-day synthesis and delivery services are available.
